# Using magnetoencephalography to track the propagation of 40 Hz invisible spectral flicker

**DOI:** 10.1101/2025.01.07.631693

**Authors:** Mark Alexander Henney, Eelke Spaak, Henrik Enggaard Hansen, Marcus Carstensen, Kristoffer Hougaard Madsen, Robert Oostenveld

## Abstract

Brain stimulation with novel 40 Hz invisible spectral flicker (ISF) has been proposed as a therapy for Alzheimer’s disease, with leading hypotheses suggesting local promotion of glymphatic clearance of amyloid as the mechanism of action. Neural signals in the gamma range and their spatial propagation over the brain can be tracked using magnetoencephalography (MEG) with high temporal and good spatial detail. However, stimulation with 40 Hz ISF requires specialised hardware which causes electromagnetic interference (EMI) with MEG equipment.

MEG measures the tiny magnetic fields of the brain, which are easily distorted by external magnetic fields. Using MEG to track the propagation of 40 Hz ISF requires multiple modifications to the experimental setup. These include, at least 1) an experimental design that promotes modulation of the neural signal, but not the artifact, 2) removal of all electronics not strictly necessary for light production from the stimulators, and 3) signal processing for 40 Hz EMI artifact suppression.

Here, we present an MEG study on the cortical propagation of 40 Hz ISF. In two experiments, we investigate the modulation that visual and non-visual cognitive tasks have on the power and propagation of cortical activity induced by 40 Hz ISF.

With the chosen experimental setup and design, the 40 Hz EMI artifact could not be entirely disentangled from the neural signal of interest, thus rendering inference on the spatial propagation of the 40 Hz signal impossible. Further improvements need to be implemented in a follow-up experimental design. We present potential solutions to allow for future investigation of 40 Hz ISF with MEG.

## 1 Introduction

### 1.1 40 Hz stimulation for Alzheimer’s

Monoclonal antibody therapy has been a major breakthrough for disease-modifying therapy of Alzheimer’s disease (AD), though their effects have been limited to a 30% reduction in disease progression [1]. Meanwhile, the incidence of amyloid-related imaging abnormalities of the oedemic or haemorrhagic subtypes has been reported upwards of 41.3% (depending on dose), of whom 26.0% experienced symptoms [2]. This benefit-to-risk ratio, combined with a price almost three times higher than the cost-effective threshold [3], motivates continuous efforts for other treatment avenues.

Notably, 40 Hz sensory brain stimulation has shown promise in pre-clinical investigations and in clinical trials of AD with positive effects reported across domains of cognition, behaviour, brain morphology, functional connectivity, and protein clearance [4–11] (see e.g., [12, 13] for overviews).

The mechanism of action for 40 Hz therapy of AD remains elusive, and the efficacy has been centre for dispute [14, 15]. A leading hypothesis for the 40 Hz stimulation modulated amyloid removal that is first described in [16] is glymphatic clearance of soluble amyloid [17], and failure thereof is considered a final common pathway to dementia [18]. Arterial vasomotor pulsation, which is entrained by the envelope of local gamma oscillations [19], regulates movement of cerebrospinal fluid (CSF) [20]. In a mouse model of AD, 40 Hz multisensory stimulation promoted cerebral vasomotion, increasing the influx of CSF and efflux of cerebral interstitial fluid to meningeal lymphatic vessels, resulting in clearance of amyloid [21]. If the benefit of 40 Hz brain stimulation is dependent on local promotion of the glymphatic flushing of soluble amyloid, it may be desirable to optimise the power and spatial propagation of the stimulus-induced activity, allthough these effort will benefit substantially from a more thorough understanding of the effects of stimulation.

Given the positive outcomes reported so far, it is worth to address the negatives, which appear to be connected to the prevailing methods of administering 40 Hz stimulation. In the OVERTURE clinical trial, using combined 40 Hz visual stimulation with luminance flicker (LF) and 40 Hz modulated 10 kHz auditory clicks led to 20.7% and 15.2% incidence of headache and tinnitus, respectively, in the active group (n=46) compared to 10.7% and 0% in the sham group (n=28) [22]. In another setting, a similar 40 Hz LF was rated by a healthy cohort for discomfort on a 0-10 Likert-type scale, scoring 6.2 higher than non-flickering continuous light [23]. This may suggest that 40 Hz LF stimulation is a suboptimal implementation of 40 Hz sensory stimulation for humans.

### 1.2 Solutions to visual stimulation side-effects

According to the Institute of Electrical and Electronics Engineers (IEEE), biological side effects of visual stimulation are associated with the perception of visual flicker [24]. Side effects may be expected from stimulation with 40 Hz LF, as a modulation depth beyond 5% is discouraged for flicker frequencies below 90 Hz [25] due to health risks [26–28]. An attractive alternative is to use a light source which, when modulated at 40 Hz, reduces or entirely removes the perception of flicker compared to LF, yet retains 40 Hz neural response. The point at which flicker becomes imperceptible, known as the critical flicker fusion frequency (CFF), is dependent on several factors of the flicker, including colour, duty cycle, wave form, and modulation depth [24].

Heterochromatic flicker (i.e., alternating between two monochromatic colours, HF) is a potential solution to reduce the CFF of therapy implementations. Optimal colour combinations have been studied, though mainly in terms of their ability to evoke steady-state potentials [29]. In an experiment with two observers of purely foveal vision, LF from a red light-emitting diode (LED) that was covering 3° of the central field-of-view (FOV) was found to fuse at ∼50 Hz [30]. The same study found that HF between the red and a luminance-matched green LED fused at ∼25 Hz. While these findings do not directly allow generalising to a larger FOV or to peripheral vision, they provide conservative lower bounds for the CFF of LF and HF, respectively. Thus, 40 Hz LF is almost guaranteed to be perceived as flickering in any implementation and thus be associated with the side effects described in [24].

Canonically, both HF and LF are associated with a high degree of perceived flicker when modulated at 40 Hz [23]. Another proposed solution is *spectral* flicker, which is a super-type of heterochromatic flicker that alternates between two *spectrally distinct light sources*. Depending on the chosen spectra, for example alternation between two spectra that both appear white, the perceived flicker can be significantly reduced while still being modulated at 40 Hz [23], thus coining the term *invisible spectral flicker* (ISF) [31].

Figure 1A schematically illustrates 40 Hz luminance flicker (LF), 40 Hz invisible spectral flicker (ISF), and non-flickering continuous (0 Hz CON) light sources on an axis of perceived flicker, with overlapping characteristics in the form of a Venn diagram.

**Fig 1.**
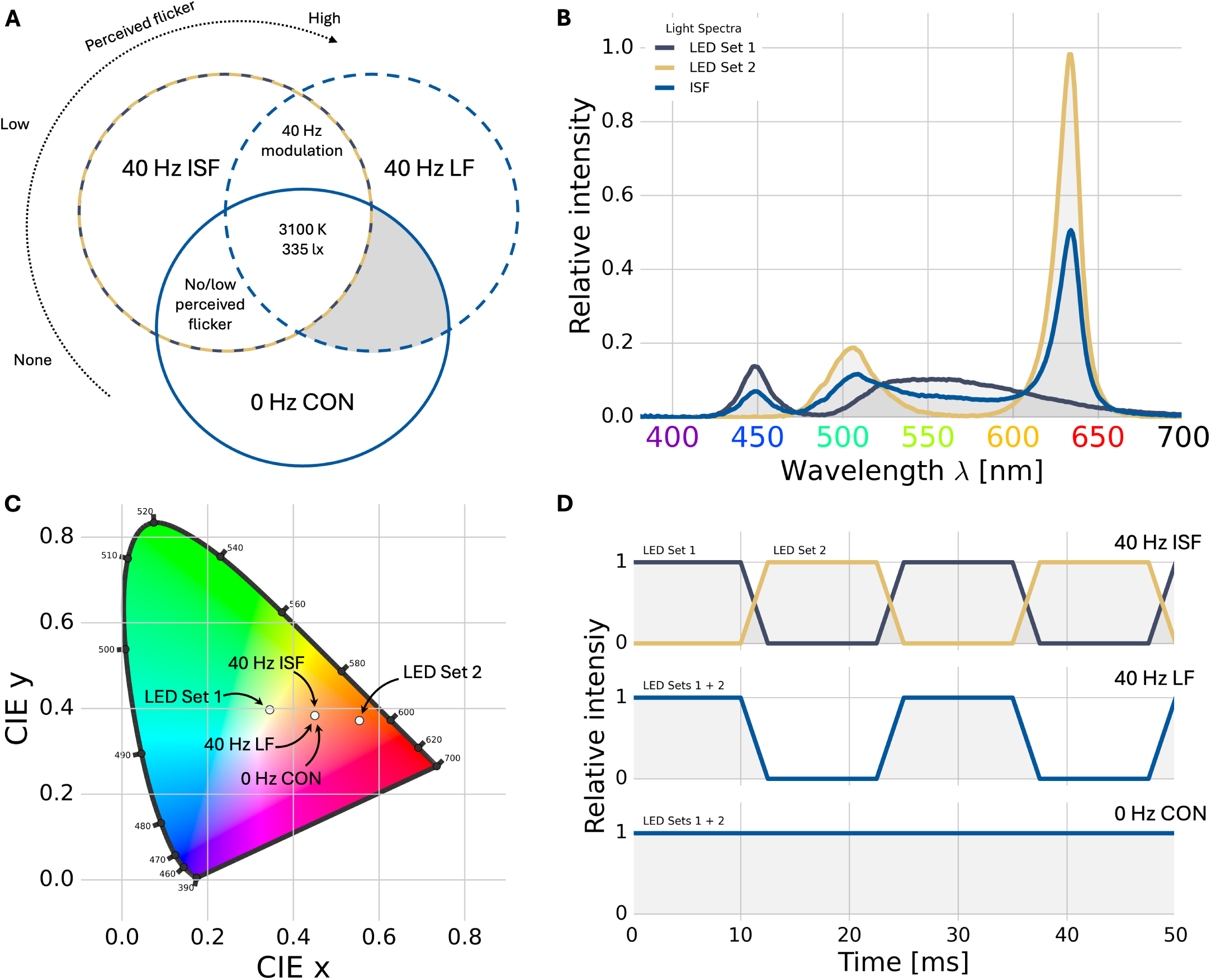
Definitions of visual stimulation paradigms: Similarities and differences between the three visual stimulation light sources, continuous light (0 Hz CON), *invisible spectral flicker* (40 Hz ISF), and luminance flicker (40 Hz LF) are summarised. **A)** Overlapping characteristics of the stimuli. All three share a correlated colour temperature of ∼3100 K and an intensity of ∼335 lx when integrated over time. The axis ”Perceived flicker” conceptualises the degree of perceived flicker, indicating that 0 Hz CON does not flicker, 40 Hz ISF has a low degree of perceived flicker, and 40 Hz LF has a high degree of flicker. **B)** The two spectra which alternate to create 40 Hz ISF, whose temporally integrated spectrum is the average of the two. Both 0 Hz CON and 40 Hz LF use the integrated ISF spectrum. **C)** The chromaticity for light-emitting diode (LED) Set 1 and LED Set 2 independently and their average, which is the perceived colour of 40 Hz ISF, 0 Hz CON and 40 Hz LF. **D)** Conceptualised modulation of the three light sources, in which 40 Hz ISF alternates between LED Set 1 and LED Set 2, 40 Hz LF alternates between the average spectrum and no light, and 0 Hz CON does not modulate.

Common to all three stimuli is their (temporally averaged) colour temperature of ∼3100 K and intensity of ∼335 lx. The 40 Hz LF and 40 Hz ISF stimuli share the 40 Hz temporal modulation, and 40 Hz ISF and 0 Hz CON share their low (or absent) perceived flicker. In fig. 1B, the two spectrally distinct phases of 40 Hz ISF (denoted LED Set 1 and LED Set 2) as well as their average spectrum (which is the basis for both 40 Hz LF and 0 Hz CON) are shown. In fig. 1C the chromaticity of each stimulus and the individual phases of 40 Hz ISF are plotted. Notably, as the 40 Hz LF and 0 Hz CON stimuli are made from the average spectrum of the 40 Hz ISF phases, the chromaticity of the three light sources are identical. Figure 1D shows the temporal modulation of each stimulus.

While the fusion of spectral flicker is very context- and viewer-dependent, and as such, the *invisibility* is circumstantial, we will use the term 40 Hz ISF to refer to the technology proposed in [31]. In [23], the perception of flicker (reported on a Likert-type scale) was highest for 40 Hz LF, followed by red/green HF, and the lowest for 40 Hz ISF. The authors also tested self-reported discomfort associated with each of the stimuli, and their ranked ordering was identical to the perceived flicker, resulting in a strong association between the two (r=0.88). They found that 40 Hz LF was perceived as more flickering and uncomfortable, compared to 40 Hz ISF.

In one task-free electroencephalography (EEG) study, 40 Hz ISF evoked about 8 dB lower occipital 40 Hz power than luminance matched 40 Hz LF [32], while in another [23] it evoked 5.4 dB lower 40 Hz global power. It is not clear how the reduction in power impacts clinical efficacy. Assuming a positive relationship between evoked power and efficacy, it is worth investigating factors that increase power while reducing discomfort or side effects.

### 1.3 Behavioural modulation of 40 Hz response

It has been proposed that the engagement in cognitive tasks during 40 Hz LF stimulation increases the power and propagation of the 40 Hz signal when compared to a task-free setting [33]. This was investigated using EEG in multiple participants and in one participant with iEEG depth electrodes that were implanted for the pre-surgical evaluation of epilepsy. In that study, participants performed a visual attention oddball experiment and a mental counting experiment, and EEG was also measured in a task-free setting. They found that, compared to the task-free setting, both the visual and the counting tasks increased the strength and extent of the 40 Hz response in EEG, and the signal propagated to deeper structures, including the hippocampus, which was not reached with task-free stimulation. These results indicate that 1) doing cognitively engaging tasks during visual stimulation improves stimulation power and propagation, and 2) the cognitive task can be visual or non-visual. Given the good temporal but limited spatial resolution of EEG, a natural next step is to investigate these factors with higher spatial resolution and to extend the design to also include 40 Hz ISF.

### 1.4 Investigating propagation

Magnetoencephalography (MEG) has superior spatial resolution and specificity compared to EEG, but also comes with specific shortcomings. One disadvantage of MEG is that the magnetic field sensors, which pick up magnetic fields around the head caused by the neuronal activity, are also sensitive to external magnetic fields. MEG recordings therefore need to be done in a magnetically shielded room (MSR). This limits the types of visual stimulation that can be delivered to participants, as the electric currents that power the light sources inside the MSR result in magnetic fields that are superimposed on the recorded signals. Typically, in steady-state visually evoked response experiments with MEG, the visual stimulation is done using a projector outside the MSR that projects through a waveguide and mirrors onto a screen in the MSR. While this is effective for presenting luminance modulation (such as LF), the colour palette of the projector is limited to combinations of only red, green, and blue (RGB) light.

The implementation of 40 Hz ISF studied previously uses four LEDs to create the two spectrally distinct colours [11, 23, 32, 34]. These spectra can not be produced with a standard RGB light source, and thus a special electronic light source is needed that is not directly electromagnetically compatible with the MEG system (see section 4.4). In a task-free functional magnetic resonance experiment of 40 Hz ISF, light was delivered by means of fibre-optic cables mounted on the head coil and fed from a customised OptoCeutics Light Therapy System (LTS; OptoCeutics ApS, Copenhagen, Denmark) [32]. While the fibre-optic cables made stimulation feasible in the task-free setting, they are impractical when the participant’s field-of-view must not be completely obstructed.

### 1.5 Research questions and feasibility

Here, we aim to overcome the incompatibility of 40 Hz ISF stimulators and MEG recording equipment through multiple avenues. We present an experimental implementation which modifies standard hardware, applies multiple steps of supervised artifact suppression, and statistically separates 40 Hz confounds from neural signals under the assumption of system linearity. The primary aim is to investigate the modulation of visual and non-visual cognitive tasks during stimulation with 40 Hz ISF on the power and propagation of the neural 40 Hz signal. As a secondary endpoint, we investigate the inverse relationship; namely the effects of different types of non-flickering, invisibly flickering, and visibly flickering lights on the performance in the cognitive tasks.

## 2 Results

In the following, we present the results of a visual attention and an arithmetic experiment implemented according to section 4 to mitigate 40 Hz EMI artifacts. First, behavioural outcomes are presented with participant response time and response accuracy. Subsequently, we present sensor-level and source-reconstructed data to address the primary research question on the modulation effects of cognitive engagement on the power and propagation of 40 Hz. Finally, we present an exploratory analysis to investigate the contribution of the stimulus artifact to the signals and the efficacy of the preprocessing and analysis procedures.

### 2.1 Visual Attention Experiment

Participants judged whether a probe stimulus was identical to or different from an earlier presented target. Before the target, they were cued to covertly attend to either the right or left side (see fig. 12 and section 4.3.1 for details).

#### 2.1.1 Behavioural response

Figure 2 presents the behavioural responses for the visual attention experiment. From panel A, it is evident that there was no difference in reaction time between the three types of visual stimulation. Panel B shows that there was also no difference in reaction time between tasks where the grating orientations were congruent or incongruent (task congruence), nor did the side of lateral attention affect the response time (panel C). The response time across conditions was distributed around 550 ms in the range of [280; 1000] ms.

**Fig 2.**
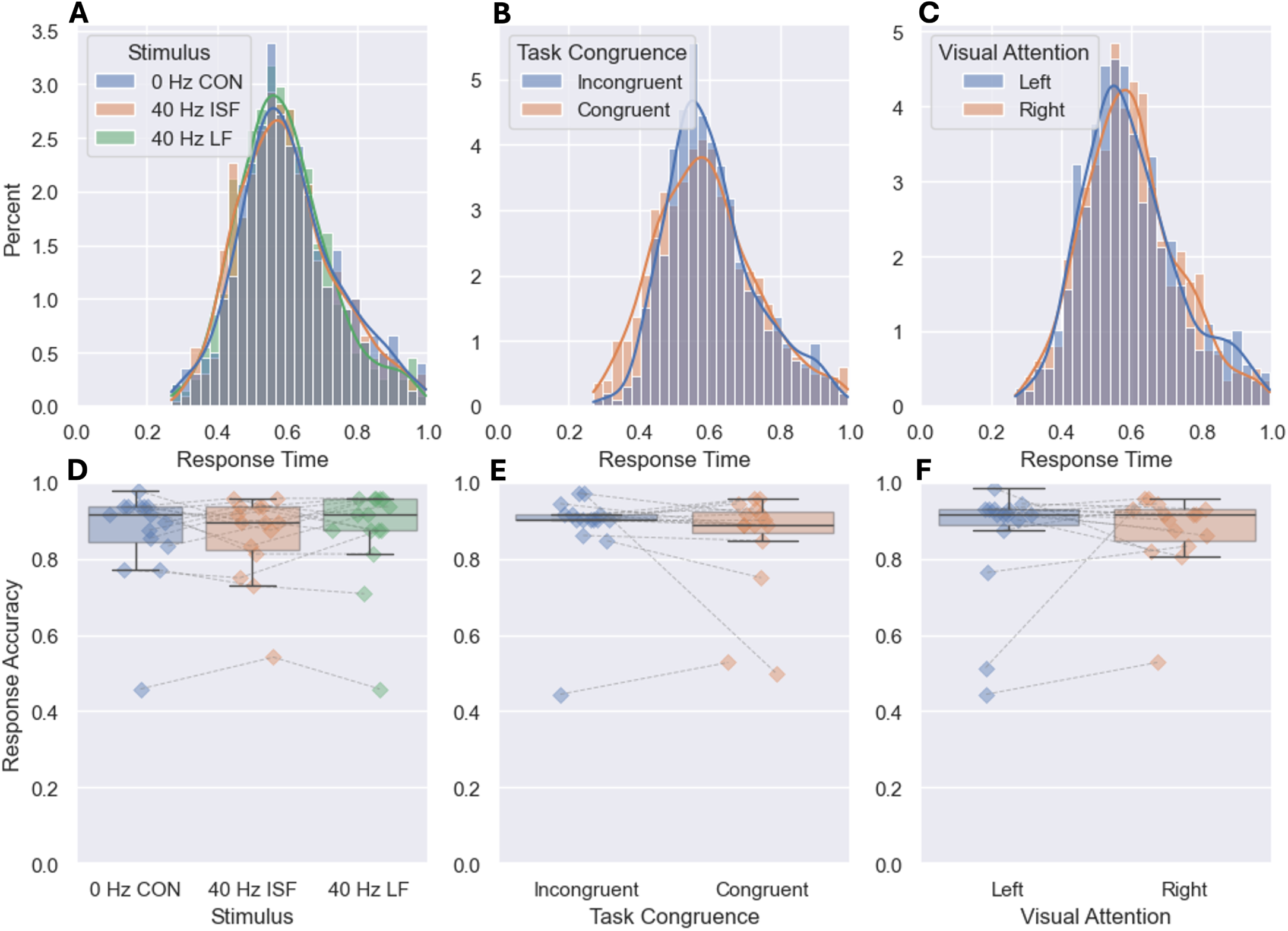
Behavioural responses for visual attention experiment: Panels **A**, **B** and **C** show reaction times, while panels **D**, **E** and **F** show the response accuracies. Both reaction time and accuracy are stratified by stimulus (left column), task congruence (middle column), and side of visual attention (right column). Connected points belong to the same participant.

The bottom row shows the response accuracy distributed around 90%, in which there was no difference between stimulation conditions (panel D), task congruence (panel E), nor side of attention (panel F). One subject consistently responded with an accuracy around 50% across conditions, suggesting that they did not fully understand the instructions and purpose of the task. Another participant responded randomly to congruent tasks and to the left attention task, also suggesting a misunderstanding of the task.

#### 2.1.2 Sensor level lateralisation of 40 Hz power

Figure 3 presents the modulation effect of lateral attention on 40 Hz power. The top row shows the grand averages of relative power spectral density between the right and left attention condition (right − left) in the [30; 50] Hz band for each channel, while the bottom row presents a topographic distribution of relative 40 Hz power between lateral conditions.

**Fig 3.**
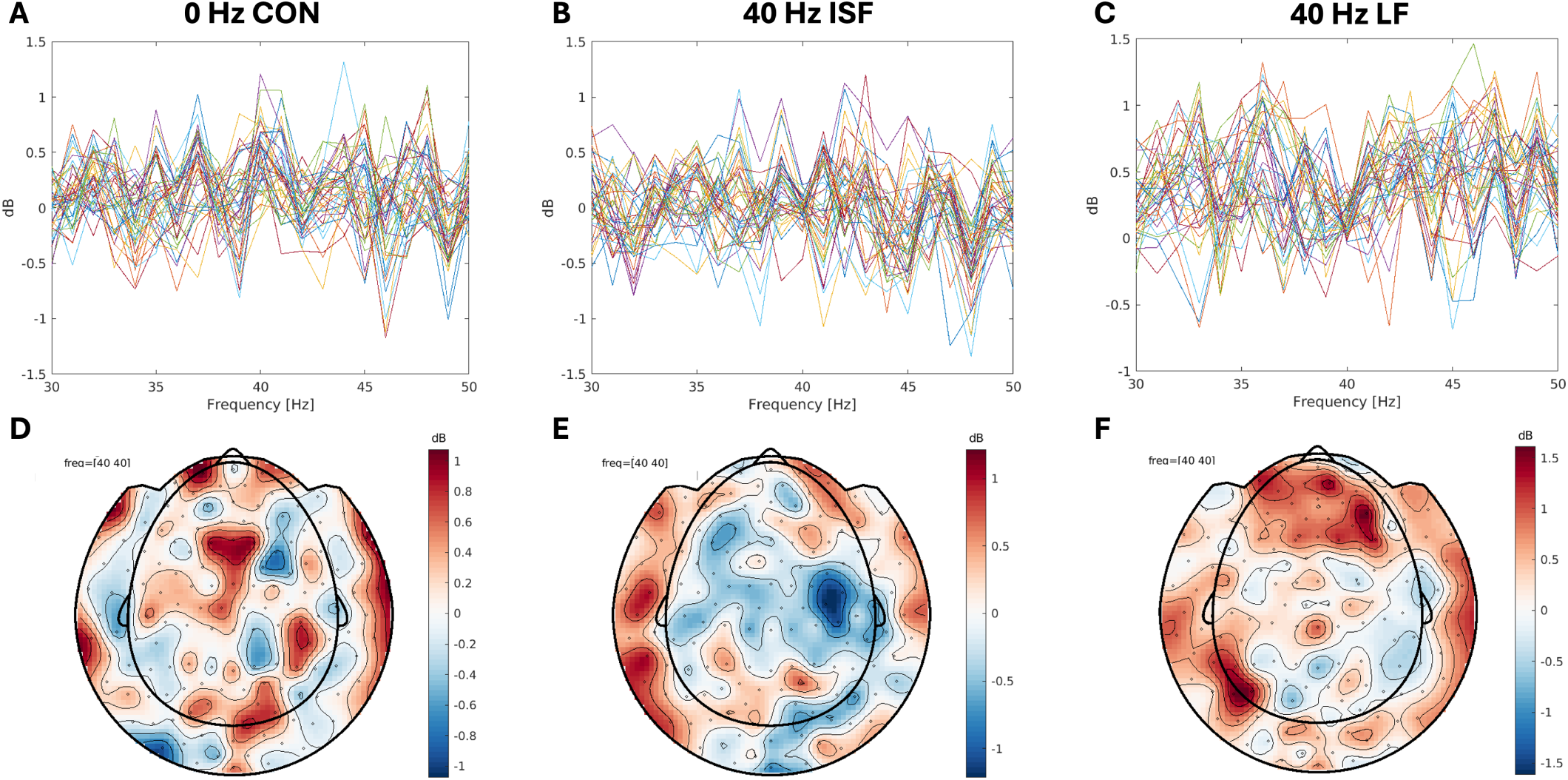
Lateral attention modulation of sensor 40 Hz power: Grand averages of the difference between right and left lateral attention conditions (right − left), stratified by stimulus. Panels **A**, **B**, and **C** show relative power in the [30; 50] Hz band for each channel for 0 Hz CON, 40 Hz ISF, and 40 Hz LF. Panels **D**, **E**, and **F** show a topographic representation of the relative 40 Hz power for 0 Hz CON, 40 Hz ISF, and 40 Hz LF.

Panels A and D in the first column show the lateral contrast between the right and left lateral attention condition during 0 Hz CON stimulus, in which the relative power is distributed around 0 dB with homogenous variance over channels over the [30; 50] Hz band, and no particular topographic pattern can be observed.

The contrast during 40 Hz ISF stimulation (panel B) reveals a similar spectral distribution around 0 dB in the [30; 50] Hz broadband, but has a notably lower variance, specifically in the 40 Hz bin. A topographic representation of the relative 40 Hz power (panel E) indicates a slight lateral asymmetry, but the difference is low, barely exceeding 1 dB.

Relative power spectra during 40 Hz LF stimulation (panel C) reveal a similar pattern to panel B, in which the [30; 50] Hz broadband displays a higher variance compared to the 40 Hz bin which has a notably lower variance. However, the power has a slight positive bias (∼0.25 dB) which is also reflected in the topography (panel F), suggesting a slightly higher 40 Hz power during the right attention condition. While there is again a slight indication of a lateral difference in the topographic pattern, a symmetric positive frontal pattern is also evident, though the relative power difference remains low.

Note that the visual stimulation was always the same in the right and left lateral attention conditions, and hence we had expected artifacts due to the stimulation to cancel out in the contrast. The variance inhomogeneity at the 40 Hz bin compared to the [30; 50] Hz broadband that is evident in the PSDs in panels B and C, but absent from panel A, indicates that the MEG system might be driven into a possible non-linear range during 40 Hz stimulation conditions. This is explored further in section 3.3.3.

#### 2.1.3 Source reconstruction

Figure 4 shows the source reconstruction of the 40 Hz signal modulated by lateral attention for each of the three types of stimuli. These are presented as statistical parametric maps (SPMs) for the group-level estimate of the right versus left lateral attention. Positive (yellow) values indicate higher power during right attention, and negative (blue) values indicate higher power during left attention.

**Fig 4.**
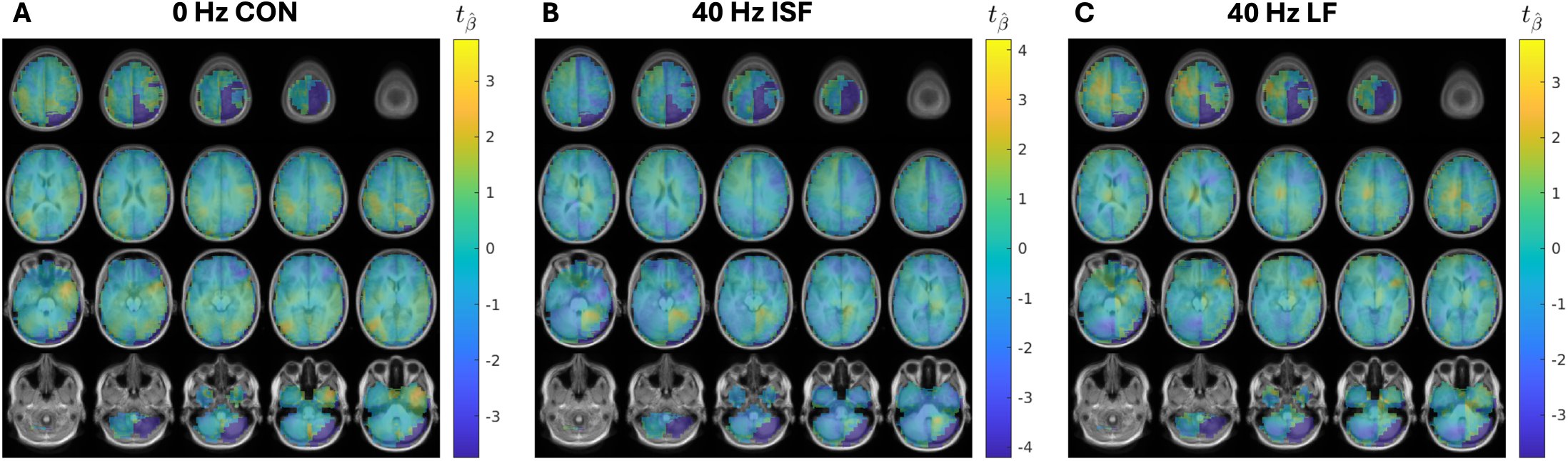
Source reconstructed visual attention modulation: Lateral (right − left) visual attention modulation of 40 Hz neural source power during **A)** 0 Hz CON stimulation, **B)** 40 Hz ISF stimulation, and **C)** 40 Hz LF stimulation. Colours represent estimated *t* -statistics for the lateral contrast at estimated sources. No significant systematic differences were established using non-parametric cluster-based permutation testing.

Panel A shows the modulation effect of lateral attention on the 40 Hz source power during 0 Hz CON stimulation. The statistical map is unmasked because the non-parametric statistical cluster-based permutation test (see section 4.5.4) revealed no significant systematic differences between the right and left attention conditions. There is no obvious pattern in the spatial distribution.

Panel B shows the modulation effect of lateral attention on the 40 Hz source power during 40 Hz ISF stimulation. As with the 0 Hz CON stimulation, the statistical test revealed no systematic differences between the attention conditions, and thus the map is unmasked. There is an indication of higher 40 Hz source power in the inferior right occipital area during right attention, but no corresponding anti-symmetric negative-valued area in the left hemisphere.

Panel C shows the modulation effect of lateral attention on the 40 Hz source power during 40 Hz LF stimulation. As with 0 Hz CON and 40 Hz ISF stimulation, the statistical test revealed no systematic differences between the attention conditions, and thus the map is unmasked. There is an indication of higher 40 Hz source power in the lower left occipital area during the left attention condition, but no opposite corresponding area in the right hemisphere. In the superior left hemisphere, the 40 Hz source power appears to be slightly higher in the right attention condition.

The source-level statistical results show no significant effects between the attention conditions and the interpretation of these results is not trivial. The lateral contrast during 0 Hz CON stimulation was designed as a negative control condition and not expected to result in systematic differences between attention conditions. Inversely, the 40 Hz LF stimulation was considered a positive control, for which an anti-symmetric lateralisation effect was expected in the the right attention versus left attention contrast. A null-finding on the positive control, in conjunction with a reduction of the variance in the signals specifically at 40 Hz described in section 3.1.2, suggests that the recovery of the 40 Hz neural signal was not sufficiently successful in the 40 Hz LF and ISF stimulation conditions. This renders any inference about the modulation effect of lateralised visual attention on the evoked 40 Hz power and propagation impossible.

### 2.2 Arithmetic Experiment

Participants judged whether the presented outcome of an earlier presented sum was correct or incorrect. The sum was either of high or low difficulty (see fig. 13 and section 4.3.2 for details).

#### 2.2.1 Behavioural response

Figure 5 shows the behavioural responses for the arithmetic experiment. Panel A shows that the response time was distributed around 550 ms in the range [280; 750] ms, and that there was no difference between the three visual stimuli. Panel B shows an effect of the task: the response time was shorter for trials where the presented outcome of the sum was correct than for trials where it was incorrect. Finally, as seen in panel C, the low arithmetic difficulty also resulted in a shorter response time than the high difficulty. In general, there was a higher occurrence of responses timing out (i.e., exceeding 750 ms) compared to the visual attention experiment.

**Fig 5.**
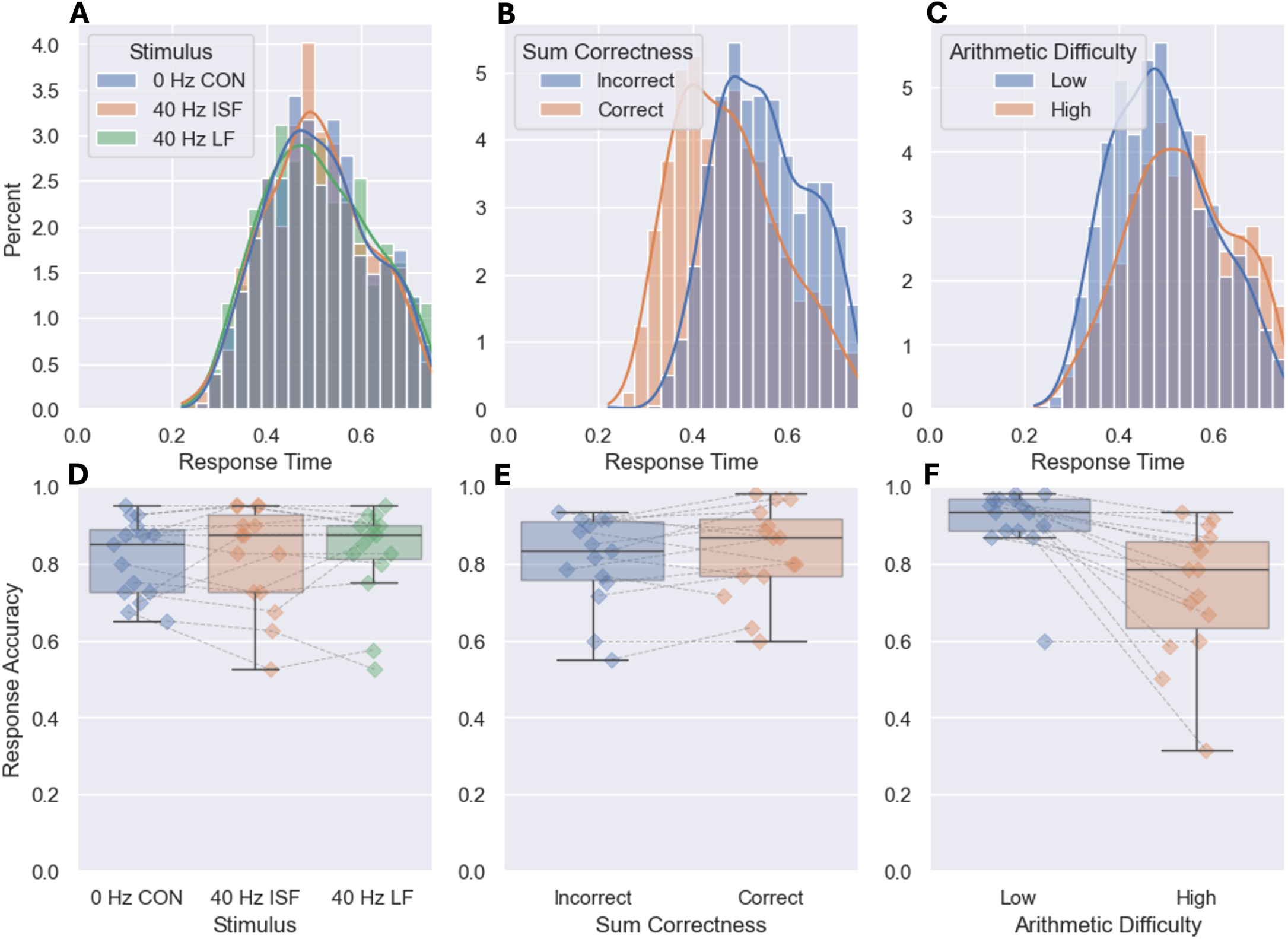
Behavioural responses for arithmetic experiment: Panels **A**, **B** and **C** show reaction times, while Panels **D**, **E** and **F** show the response accuracy. Both are stratified by stimulus (left column), sum correctness (middle column), and difficulty of arithmetic (right column). Connected points belong to the same participant.

The response accuracies were approximately the same across the three visual stimuli (panel D). There was a slight indication that participants responded correctly more often when the result of the sum was correct, than when it was not (panel E), though the difference is small. There was a big difference of approximately 20% in the response accuracy between tasks with low and high arithmetic difficulty (panel F). The latter suggests that the difficulty levels were chosen appropriately.

#### 2.2.2 Sensor level 40 Hz power

Figure 6 presents the modulation effect of arithmetic difficulty on 40 Hz power. The top row shows the grand averages of relative power spectral density between the high and low arithmetic difficulty conditions (high − low) in the [30; 50] Hz band for each channel, while the bottom row presents topographic distributions of relative 40 Hz power between arithmetic conditions.

**Fig 6.**
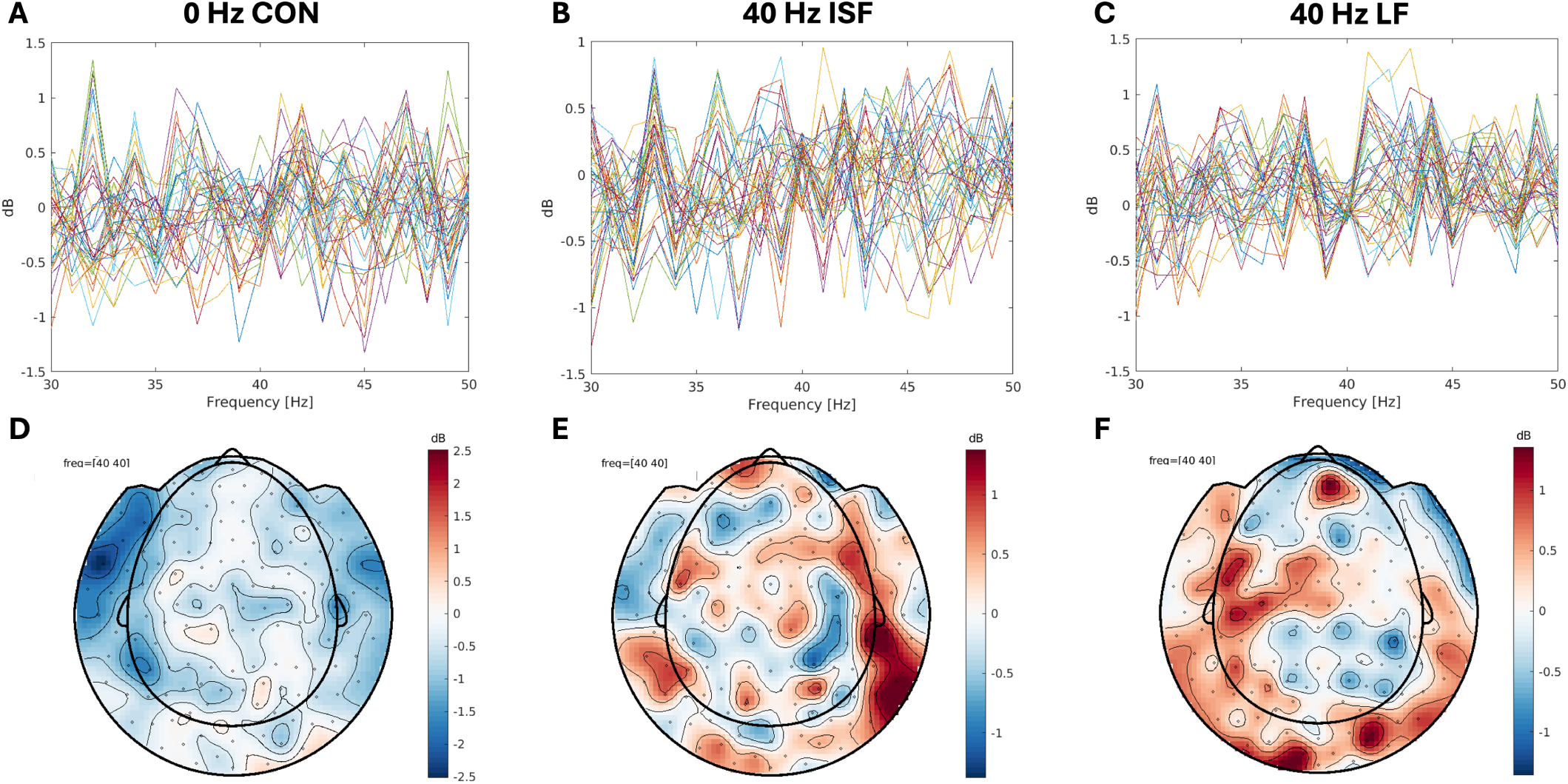
Arithmetic difficulty modulation sensor 40 Hz power: Grand averages of the difference between high and low difficulty conditions, stratified by stimulation type. Panels **A**, **B** and **C** show relative power in the [30; 50] Hz band for each channel for 0 Hz CON, 40 Hz ISF, and 40 Hz LF. Panels **D**, **E** and **F** show topographic representations of the relative 40 Hz power for 0 Hz CON, 40 Hz ISF, and 40 Hz LF.

Panels A and D in the first column show the high versus low arithmetic difficulty contrast during 0 Hz CON stimulus. The relative power is distributed around 0 dB with homogenous variance between channels over the [30; 50] Hz band, and no particular topographic pattern is seen.

The contrast during 40 Hz ISF stimulation (panel B) reveals a distribution around 0 dB in the [30; 50] Hz broadband, but a notably lower variance at 40 Hz. This indicates, again, a non-linear phenomenon specifically in the 40 Hz bin, similar to the lateral attention experiment (explored further in section 3.3.3). The topographic representation of the relative 40 Hz power (panel E) shows no particular pattern.

The relative power during 40 Hz LF stimulation (panel C) is similar to the 40 Hz ISF condition, and the non-linear phenomenon at 40 Hz is even more pronounced with a clear notch in the spectrum. The topographic representation of the relative 40 Hz power (panel F) potentially indicates a slightly higher occipital and frontal power in the high difficulty setting, as well as a slight central asymmetry.

#### 2.2.3 Source reconstruction

Figure 7 shows the source reconstruction of the 40 Hz signal modulated by arithmetic difficulty for each of the three stimuli types. They are presented as SPMs for the group level estimate of the high versus low arithmetic difficulty. Positive (yellow) values indicate higher power during the high-difficulty condition, whereas negative (blue) values indicate higher power during the low-difficulty condition.

**Fig 7.**
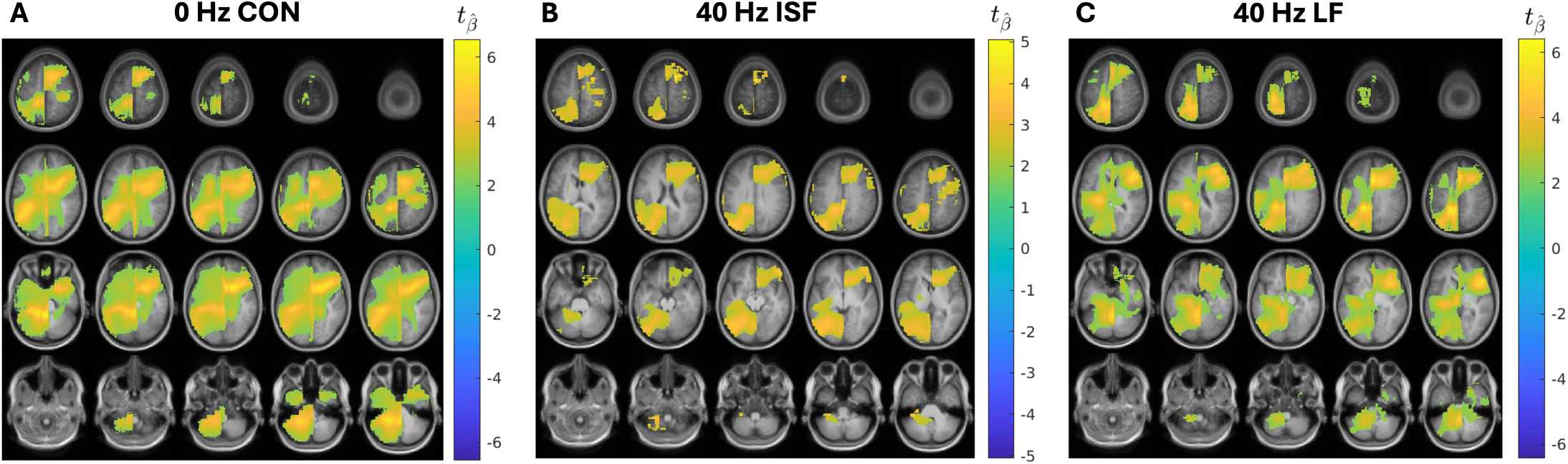
Source reconstructed arithmetic difficulty modulation: Arithmetic difficulty (high − low) modulation of 40 Hz neural source power during **A** 0 Hz CON stimulation, **B** 40 Hz ISF stimulation, and **C** 40 Hz LF stimulation. Colours represent estimated *t* -statistics for the arithmetic difficulty contrast at estimated sources. Values are masked by systematic significances established using non-parametric cluster-based permutation testing.

Panel A shows the modulation effect of higher arithmetic difficulty on the 40 Hz source power during 0 Hz CON stimulation. The SPM is masked to only include systematic differences between the high- and low arithmetic difficulty, established using a non-parametric cluster-based permutation test (see section 4.5.4). We find a cluster with widespread support, indicating that there was systematically higher 40 Hz power during high compared to low arithmetic difficulty. The pattern, however, is asymmetric with power differences peaking more anteriorly in the right hemisphere than in the left hemisphere.

Panel B shows a similar map of the modulation of the arithmetic difficulty on the 40 Hz source power during 40 Hz ISF stimulation, masked by significance. There are two large clusters, suggesting a higher 40 Hz power in the right anterior and left posterior areas during high arithmetic difficulty. The support is narrower than in the 0 Hz CON condition, but the pattern is similar.

Panel C shows the modulation of the arithmetic difficulty on the 40 Hz source power during 40 Hz LF stimulation, masked by significance. The pattern, again, is one large asymmetric cluster with support in the anterior right area and the posterior left area.

As with the visual attention experiment, the results do not allow for an easy interpretation. While there is a systematic difference between high and low arithmetic difficulty in terms of 40 Hz power and propagation for all three flicker conditions, we cannot provide a neurophysiological explanation for them. A significant modulation of 40 Hz source power in the control condition with 0 Hz CON stimulation, for which there is no input 40 Hz signal to modulate, would indicate that the change is a broad-band effect. However, the sensor level analysis (see section 3.2.2) found the relative sensor power in the [30; 50] Hz band to be distributed around 0 dB, which does not corroborate that notion. In addition, it is unexpected that the 40 Hz LF positive control condition would result in an asymmetric and almost identical pattern to the negative control. Considering these findings, it is unlikely that the significant clusters reflect neurophysiological difference. Rather, they may be the product of imperfect elimination of stimulation hardware artifacts, even by spatial filtering with the beamformer.

### 2.3 Exploratory validation results

Given that both the sensor level spectral analyses and beamformer source reconstruction of the 40 Hz neural signals were unsuccessful, we present further exploratory results here. These are intended to verify to what extent our artifact correction procedures were successful, or not, in order to guide future work.

#### 2.3.1 Artifact removal

The result of the de-noising procedure (see section 4.5.1) is presented in fig. 8 for a random sample of 10 trials with 40 Hz stimulation from one representative channel and subject during the visual attention experiment.

**Fig 8.**
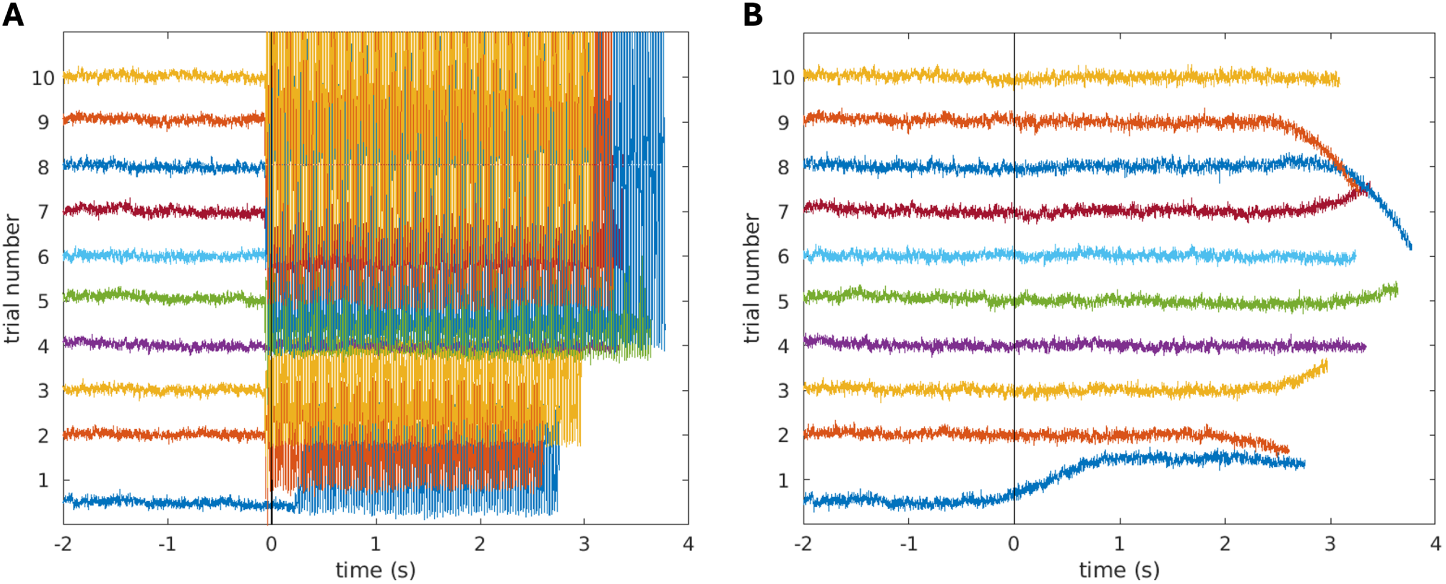
Artifact removal by linear transformations: The 40 Hz electromagnetic interference (EMI) artifact is actively suppressed by means of 3rd order gradient rebalancing and subsequent identification of residual 40 Hz power in the reference gradiometers with principal component analysis (PCA), which was projected and subtracted from the sensors. Panel **A** shows 10 representative trials with the 40 Hz EMI artifact present from the stimulus onset at *t* = 0 s. In panel **B** the artifact appears to have been removed using third-order synthetic gradiometer rebalancing and subsequent PCA denoising.

Panel A shows the time series aligned to the stimulus onset at 0 s, at which point the 40 Hz EMI artifact begins. Panel B shows how the artifact appears to have been successfully removed. This visually confirms that the noise suppression effectively reduced the variance of the 40 Hz EMI artifact.

#### 2.3.2 Confirming sensor level 40 Hz presence

While fig. 8 visually confirms a reduction in variance associated with the visual stimulators, we must also verify that there is a 40 Hz signal present during 40 Hz ISF stimulation, and none during 0 Hz CON, to ensure that neural signals were not entirely suppressed in the process. Figure 9 shows three representative grand-averaged PSDs taken from the visual attention experiment to address this concern.

**Fig 9.**
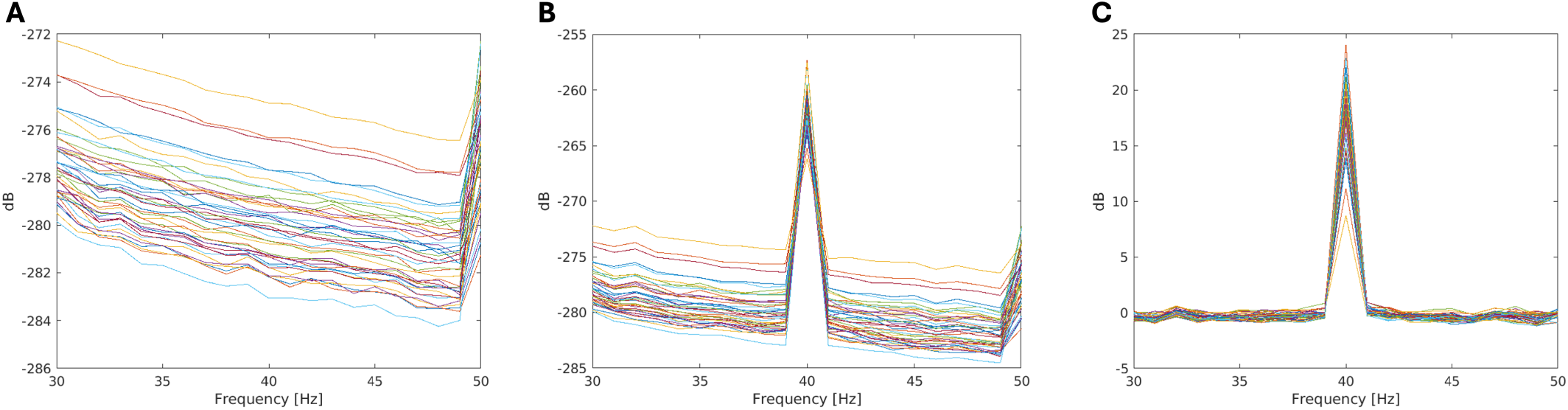
Verifying the preservation of 40 Hz power: To ensure that not all 40 Hz power had been suppressed by the 40 Hz electromagnetic interference artifact removal procedure, the average power spectral densities (PSDs) for 0 Hz CON and 40 Hz ISF and the difference between them are plotted. **A)** The PSD for 0 Hz CON shows an expected aperiodic 1*/f ^a^* shape with only a periodic 50 Hz line noise peak. **B)** The PSD for 40 Hz ISF has a clear 40 Hz peak ∼15 dB higher than the 50 Hz line noise peak. **C)** The difference in PSD between 0 Hz CON and 40 Hz ISF is 0 dB throughout the [30; 50] Hz band with the exception of the 40 Hz peak of [10; 25] dB height.

Panel A of fig. 9 shows the power in the [30; 50] Hz band during 0 Hz CON stimulation, in which we expect an entirely aperiodic 1*/f^a^* shape, with the exception of the 50 Hz line noise peak. This confirms that 40 Hz power has neither crept in, nor has been suppressed by the linear transformation while the 0 Hz CON stimulation was active.

Panel B of fig. 9 shows the power in the [30; 50] Hz band during 40 Hz ISF stimulation, in which we expect an aperiodic 1*/f^a^* component with periodic 50 Hz line noise identical to the 0 Hz CON, but also with a periodic 40 Hz signal superimposed thereon. This appears to be the case, confirming that a 40 Hz signal from the brain remains after artifact removal. Panel C shows the relative power between 0 Hz CON and 40 Hz ISF, in which the 40 Hz peak should be the only difference, which is also the case.

#### 2.3.3 Suspected non-linearities

From the PSDs in fig. 9 in conjunction with the time series in fig. 8, there are no indications that the artifact removal was unsuccessful. However, as evident from the spectral analysis of the behavioural contrasts within stimulus types (see figs. 3 and 6), the stimulators may have driven the sensors into a non-linear range. This is seen in fig. 10 as a notch, i.e., a reduction in the variance of power over channels, specifically at 40 Hz and only for the 40 Hz ISF and LF stimulus conditions. This renders the planned elimination of residual artifacts by conditional contrasting impossible, leaving an unresolvable statistical confound in the data.

**Fig 10.**
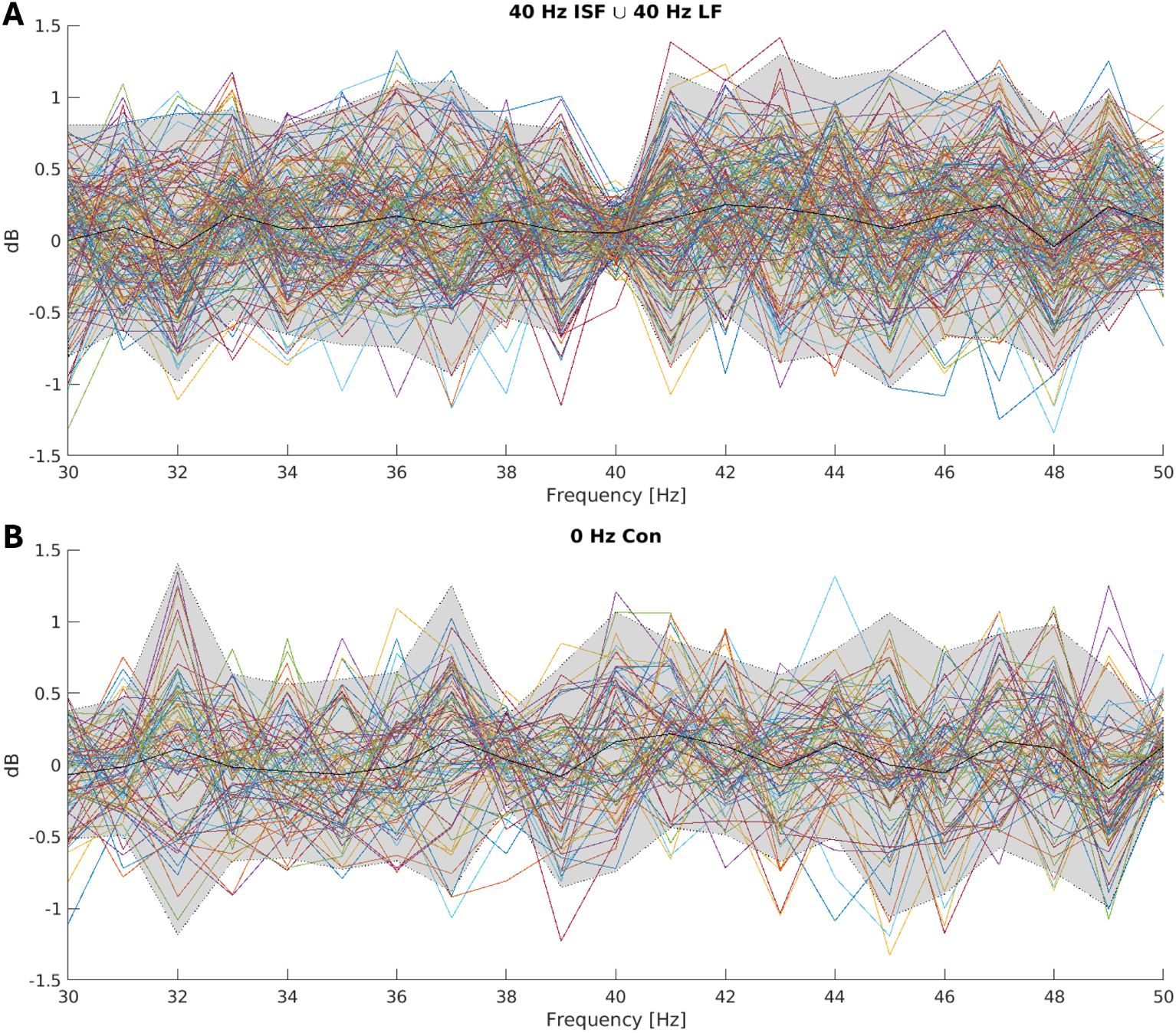
Suspected non-linearity in sensors: Power spectral density is plotted across sensors with shaded area showing the variance (scaled by a factor five for visualisation). **A)** The variance is markedly reduced at specifically 40 Hz (pooled for 40 Hz ISF and LF stimulus conditions). **B)** In comparison, the variance is homogenous over frequencies for the 0 Hz CON stimulus condition.

## 3 Materials and methods

### 3.1 Participants and ethics

The experiment was proposed to and approved by the institutional review board at the Donders Institute for Brain, Cognition, and Behaviour, Radboud University, The Netherlands, as part of the protocol ’Imaging Human Cognition’ (NL45659.091.14), which was approved by METC Oost-Nederland (2014/288) in accordance with the Declaration of Helsinki. Fifteen healthy adult volunteers were recruited from the Radboud University participant pool and included in the experiment after receiving written and verbal information and providing written consent. Data was collected at the Donders Centre for Cognitive Neuroimaging between April 17th and May 7th of 2024.

All participants had normal vision or could verifiably complete the cognitive tasks without the use of corrective glasses. The exclusion criteria were light sensitivity, migraine, photosensitive epilepsy, the presence of a dental wire, or any known (history of) neurological or psychiatric illness.

### 3.2 Data acquisition

Data was collected in a magnetically shielded room (MSR) using a 275-channel superconducting quantum interference device (SQUID) based MEG system (CTF MEG Neuro Innovations Inc., Coquitlam, BC, Canada) in the sitting position. Task instructions were displayed to the participants on a projection screen situated ∼70 cm in front of them, using the regular projector located outside the MSR. Visual stimulators (see section 4.4) were placed on either side tangential to the lower corners of the screens (see fig. 11).

**Fig 11.**
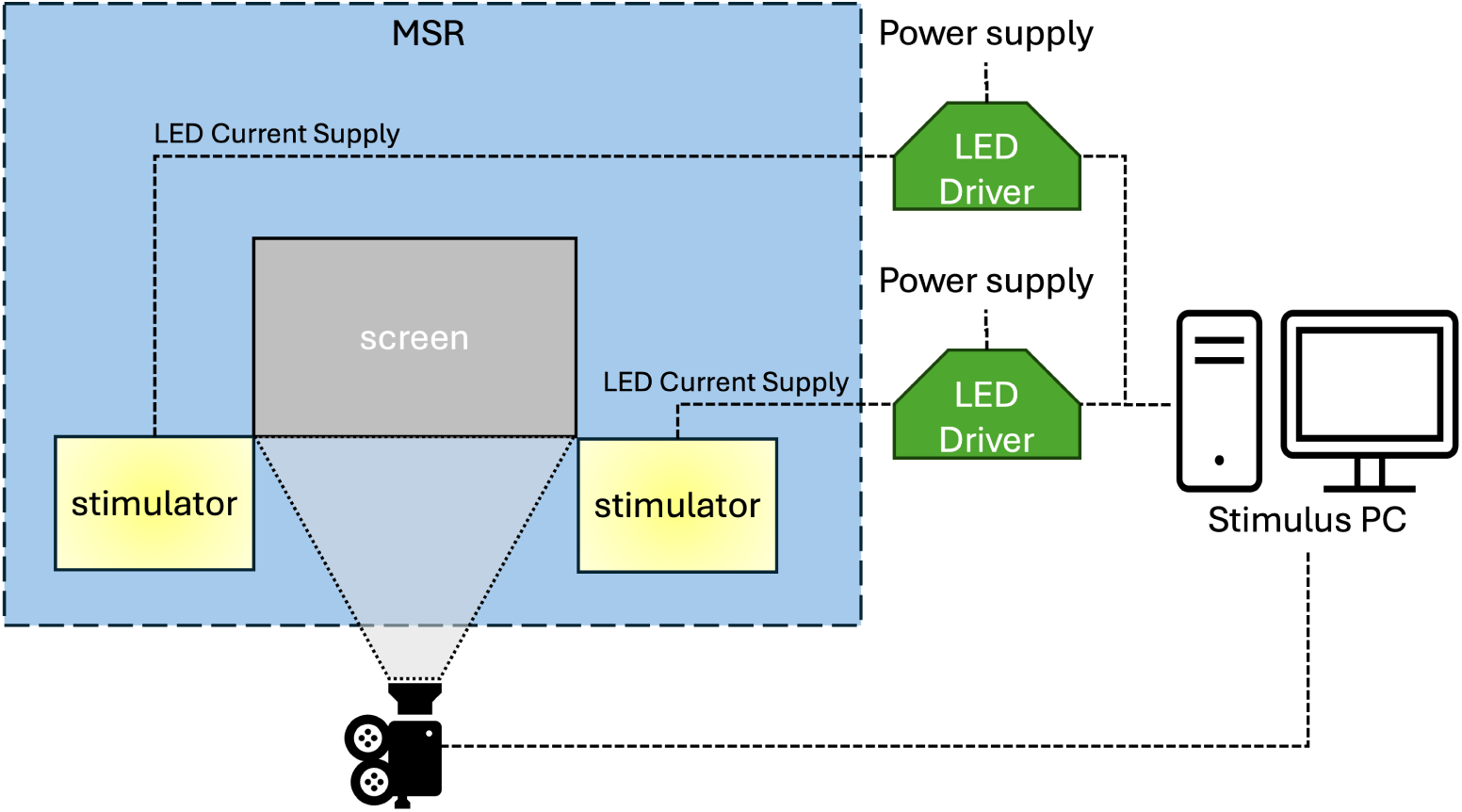
Setup of devices in magnetically shielded room: Printed circuit boards (PCBs) contain the driver electronics for the LED chips. These are powered by 12 V, 3 A power supplies outside the magnetically shielded room (MSR) and deliver current via a cable of twisted pairs to the LED chips (see section 4.4) placed inside the MSR. The stimulus PC simultaneously controls the stimulation devices and the screen display.

The experiments were controlled using a custom Python 3.8 module, developed for the project, which interfaced simultaneously with the presentation screen in the MSR using a modified PsychoPy version 2023.2.3, the stimulators using a proprietary application programming interface provided by OptoCeutics, and the event triggers integrated with the CTF system via serial connection.

### 3.3 Experimental design

To resolve the confound of stimulator artifacts, both cognitive tasks were designed with two levels orthogonalised relative to the stimulus conditions. Assuming the stimulus artifact to be independent from the task, and thus constant between levels, the contrast between task levels would eliminate the artifact confounding the 40 Hz neural signal and reveal only the task effect as event-related lateralised fields [35]. Three visual stimulation conditions, delivered by two MEG-customised visual stimulator devices (see section 4.4 and fig. 14), were used: 0 Hz CON, 40 Hz ISF and 40 Hz LF. The relationship between the three stimuli on the two axes is illustrated in fig. 1. With these, we aim to answer three questions: 1) can MEG be used to track the propagation of 40 Hz ISF? 2) does cognitive load during 40 Hz visual stimulation affect the power and/or propagation of the 40 Hz signal, and 3) is the power and propagation dependent on the perceptibility of the flicker?

**Fig 12.**
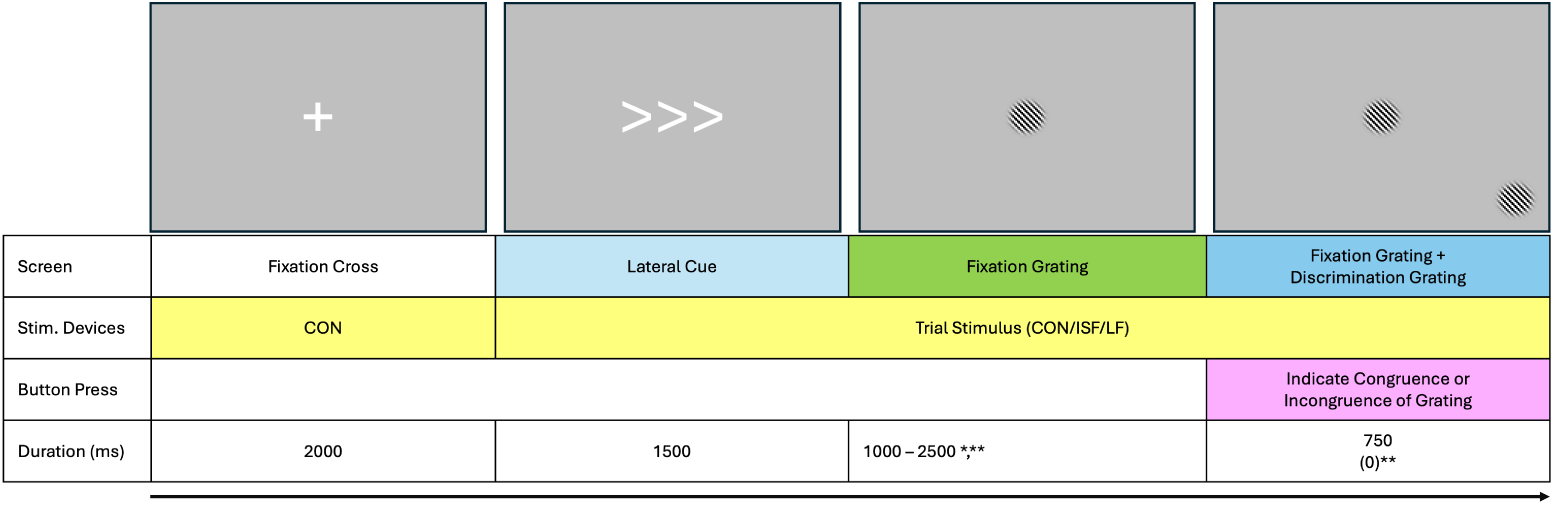
Visual attention experiment: Each trial begins with 2 s where the stimulators are on the 0 Hz CON setting and the presentation screen showing a fixation cross. Then, the stimulators switch to the stimulus setting for the given trial (0 Hz CON, 40 Hz ISF, or 40 Hz LF), and for 1.5 s, the presentation screen shows a lateral cue in the form of arrows to the left or right. Then, the arrow cues are replaced with a grating in the centre, termed the ”fixation grating”, which is oriented at a 45° diagonal and remains for a variable duration of 1 s to 2.5 s . (*) With 10% probability, the fixation grating will show for only 500 – 1000 ms (”quick trial”). Finally, a second ”discrimination grating” is shown for 0.75 s in the bottom corner (next to the stimulator) on the side which was cued. Its orientation is either the same as the fixation grating (congruent) or opposite (incongruent). During this time, the participant must indicate by button press whether the orientations are congruent or not. (**) Given the fixation grating duration exceeds 1000 ms, there is 10 % probability the fixation grating lasts the full 2500 ms, and the discrimination grating never shows (”catch trial”). All durations were sampled from a uniform distribution, and all random draws were made at runtime.

**Fig 13.**
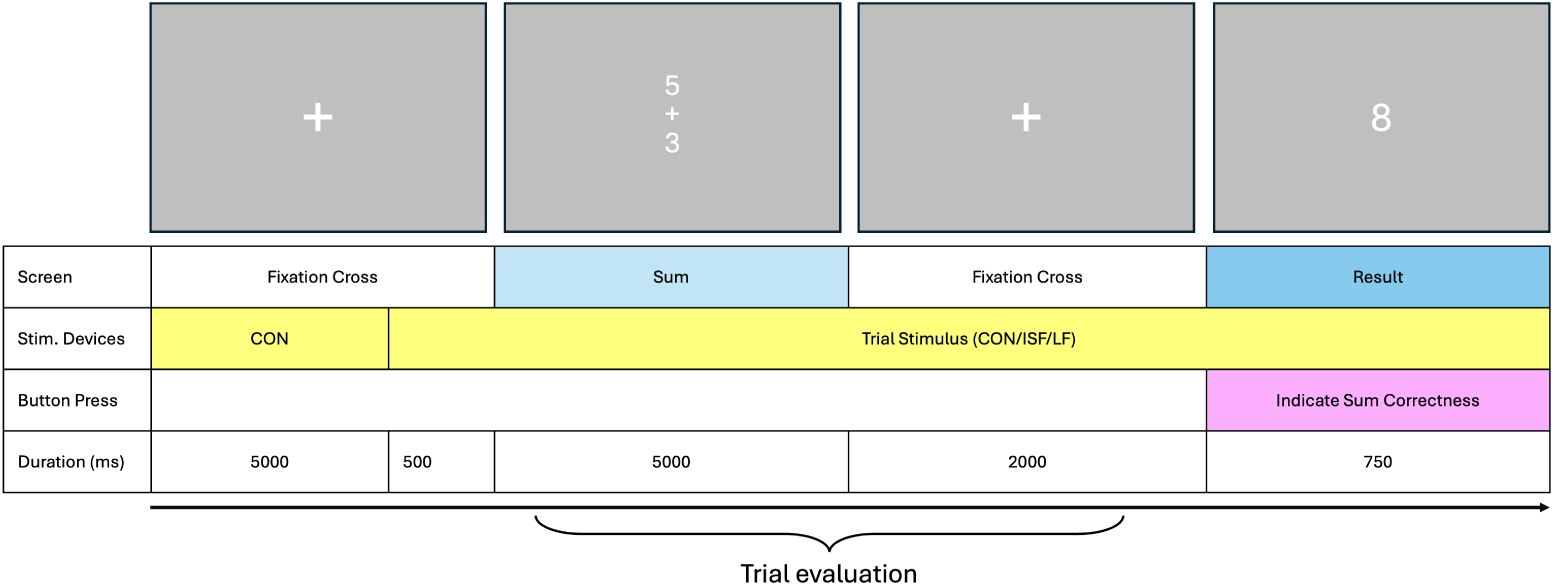
Arithmetic experiment: Each trial begins with 5 s where the stimulators are on the 0 Hz CON setting and the presentation screen showing a fixation cross. Subsequently, the stimulators switch to the stimulus setting for the given trial (0 Hz CON, 40 Hz ISF, or 40 Hz LF), while the fixation cross remains for another 0.5 s to accommodate transient effects of stimulus onset. Then, either an easy or a difficult sum is presented for 5 s, after which the fixation cross returns for 2 s. Finally, the outcome of the sum is presented for 0.75 s, which can be either correct or incorrect. During this period, the participant has to indicate by a button press, whether the presented outcome is correct.

**Fig 14.**
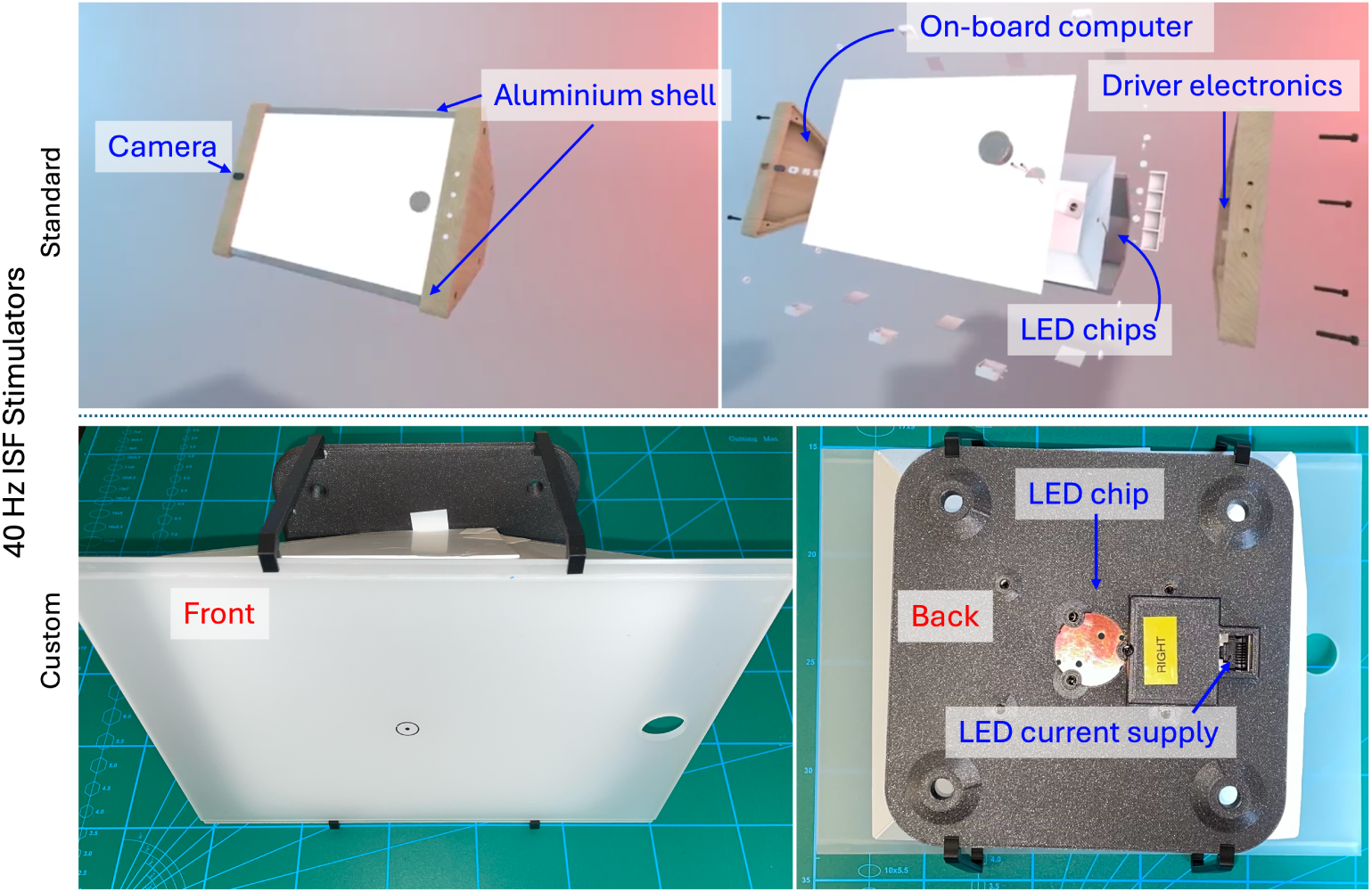
Customisation of visual stimulation device: Some components of the standard 40 Hz ISF stimulator are incompatible with the MEG system due to electromagnetic emissions.

#### 3.3.1 Visual attention experiment

For the visual task, the difference in cognitive load was implemented as a lateralised attention task with a contrast between left and right attention. The experiment (see fig. 12) started with a fixation cross shown on the presentation screen for 2000 ms, followed by an arrow cue indicating that participants should attend either left or right for 1500 ms.

Simultaneously with presenting the cue for the attention task, both the stimulators changed from 0 Hz CON light to the trial specific stimulus (either 0 Hz CON, 40 Hz ISF, or 40 Hz LF). From this time on, participants were overtly attending towards the left or right direction.

Following the cue for the attention direction, a grating (”fixation grating”) was shown for 1000–2500 ms in the centre of the screen for the subject to maintain their central gaze while they attended to the cued side. In 5% of the trials the period was only 500–1000 ms (”quick trials”) as to break expectations for the lower bound on the duration; these trials were discarded in the subsequent analysis.

Subsequently, a second grating (”discrimination grating”) was shown in the lower corner on the side that had been cued for up to 750 ms, and the orientation would be either congruent or incongruent to that of the fixation grating. When the discrimination grating was shown, subjects had to respond as quickly as possible with a button press whether the orientations were congruent or incongruent. In 5% of the trials (”catch trial”) no discrimination grating was shown to break expectations for the upper bound of the waiting duration.

Trials were evaluated in a fixed 2000 ms window starting from 500 ms after the lateral cue, thereby eliminating transients from the stimulus changing. After responding by a button press or after exceeding the 750 ms response period, the fixation cross returned, and the stimulators were changed back to 0 Hz CON.

The lateralisation sides, congruence, and stimulus factors were randomised and balanced, and all combinations were repeated twice within each of six blocks, making it a randomised complete block design (RCBD), not accounting for the independent and random selection of quick and catch trials at run time.

#### 3.3.2 Arithmetic experiment

The non-visual task was implemented as an arithmetic experiment with two levels of difficulty. The experiment (see fig. 13) started with a fixation cross shown on the presentation screen for a duration of 5500 ms to; this was longer than in the visual task, to accommodate the need for more resting time between the cognitively strenuous arithmetic.

After 5000 ms, both the stimulators changed from 0 Hz CON light to the trial specific stimulus (0 Hz CON; 40 Hz ISF; 40 Hz LF) to allow for a 500 ms transient period before showing a sum. The sum would be either low difficulty (each term *x*_low_ ∈ N: *x*_low_ ≤ 9) or high difficulty (each term *x*_high_ ∈ N: 100 ≤ *x*_high_ ≤ 400) and was shown for 5000 ms.

Subsequently, a 2000 ms interim period ensued, were the sum was replaced by a fixation cross. Finally, an outcome for the sum was shown that was either correct or incorrect. When the sum was incorrect, it was offset by Δ*x*_low_ ∈ {1, 2} in the low difficulty level and by Δ*x*_high_ ∈ N: 10 ≤ Δ*x*_high_ ≤ 20 in the high difficulty level. The result was shown for up to 750 ms, during which the subject had to indicate by button press whether the presented outcome was correct or not.

Trials were evaluated in a fixed 6000 ms window starting from the point of sum presentation. After responding by a button press or after exceeding the 750 ms response period, the fixation cross returned, and the stimulators were changed back to 0 Hz CON.

The design, again, was an RCBD with difficulty levels, sum correctness, and stimulus factors randomised and balanced in all combinations of levels within each of 10 blocks.

### 3.4 Visual stimulation devices

The visual stimulation devices were customised versions of the Light Therapy System (LTS; OptoCeutics ApS, Copenhagen, Denmark), adapted to improve electromagnetic compatibility and deliver three types of visual stimuli: 0 Hz CON, 40 Hz ISF, and 40 Hz LF.

To restate, the two independent white spectra in the modulated 40 Hz ISF light rules out standard RGB light sources, and thus the necessity of using the specific system rather than standard displays or projectors. However, this system was not designed to operate in experimental environments sensitive to electromagnetic interference. Precautions therefore had to be taken with regards to emissions from not only driving the LEDs, but also with the periodic modulation of the LEDs necessary for the stimulation, see fig. 1, panel D on page 4.

To overcome these issues, customisations were made to the devices. This included a decoupling of the stimulator (LEDs) and the electromagnetically noisy components (LED drivers and control circuitry). The resulting stimulators housed the LEDs with a polyethylene reflector and an acrylic diffuser (see fig. 14). The two stimulators were positioned adjacent to the lower corners of the presentation screen in the MSR, and the LED chips were connected by twisted-pair cables to the driver electronics and power supplies outside the MSR. Both of the driver electronics boards were controlled by the stimulus computer (see fig. 11).

The LED colour profiles were centred (± full-width at half maximum) around, blue (460 ± 30 nm), cyan (500 ± 30 nm), lime (555 ± 110 nm), and red (630 ± 30 nm). The brightness and waveforms were controlled by pulse-width modulation (PWM) at 2.93 kHz. In the standard 40 Hz ISF stimulator configuration, there are two LED chips, and the forward driving currents to each of them are fixed at 500-900 mA, depending on the LED. To prospectively reduce the 40 Hz EMI while maintaining high illumination, the custom stimulators used only one LED chip each, thereby halving the voltage, and the currents were minimised to 181-370 mA by increasing the PWM on-time to the maximal value within the linear region.

The two devices produced equivalent light according to the 40 Hz ISF light specifications of correlated colour temperature (CCT) and average spectral intensity, integrated over an entire 25 ms cycle. This was achieved at CCTs of 3153 K and 3122 K and at intensities of 337 lx and 334 lx at 30 cm, respectively. The modulation depths between the two spectra in the 40 Hz ISF were 4.78% and 1.93%, respectively, which is within the accepted tolerance of 5%.

### 3.5 Data analysis

Data analysis was done with MATLAB 2023b (Mathworks, Nattick, MA) using a forked and modified version of the FieldTrip toolbox [36] and custom analysis scripts.

These include primarily the on-board computer and LED driver electronics. To improve electromagnetic compatibility, the LED drivers and their power supplies were moved out of the MSR according to fig. 11 with currents delivered over twisted-pair cables into the MSR for the LED chips. **Top row:** Standard 40 Hz ISF stimulator with problematic components annotated with blue. **Bottom row:** MEG-customised 40 Hz ISF stimulator with only the LED component and connector for current supply.

#### 3.5.1 Data cleaning

For each MEG recording, we applied two steps of linear data transformations to remove the 40 Hz EMI artifact. First, we applied third-order synthetic gradient rebalancing [37] with FieldTrip’s ft denoise synthetic. Subsequently, we used principal component analysis (PCA) to identify that most variance in the reference gradiometers could be explained by two principal components. Under the assumption that these components made up the 40 Hz EMI artifact, we employed FieldTrip’s ft denoise pca to project them onto the data space and remove them from the signal.

#### 3.5.2 Spectral analysis

Within each trial and per channel, the power spectral density (PSD) was estimated as a periodogram:

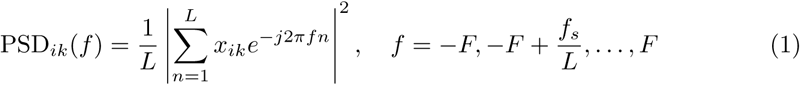

for trial *i*, channel *k*, sampling frequency *f_s_*= 1200, *F* = *^fs^*, and *L* is the duration of the period of interest (defined in sections 4.3.1 and 4.3.2).

Within subjects and for each channel, the absolute 40 Hz power was averaged for each stimulus and task condition in the experimental design. Contrasts between conditions were computed based on the averages and brought forward for group-level analysis.

For topographic presentation of the sensor level 40 Hz power, averaged data was transformed to a planar gradient configuration using FieldTrip’s ft megplanar, and the magnitude was estimated using ft combineplanar.

#### 3.5.3 Source reconstruction

We reconstructed the neural sources using spatial filtering by linearly constrained minimum variance (LCMV) beamforming [38]. Beamforming not only localises the sources of the MEG signal, but also suppresses interfering contributions from non-brain sources, thereby further contributing to noise suppression [39].

The co-registration between MEG and MRI spaces was done using head-localizer coils that were placed at the nasion and on ear molds placed in the left and right ear. A T1-weighted gradient echo sequence was used to acquire the anatomical MRI. Lead fields were estimated based on a single-shell volume conduction head model [40] which had been non-linearly warped from a template brain to the individuals head shape [41]. The source model was defined on a three-dimensional grid in one hemisphere with equidistant spacing of 8 mm with a symmetry constraint for the LCMV beamformer to fit double-dipoles [42]. This was done in anticipation of symmetrically correlated but spatially distinct sources from the steady-state evoked responses to bi-lateral stimulation which are attenuated and spatially distorted by the LCMV algorithm [38]. Prior to beamformer source reconstruction, the artifact-removed data was band-pass filtered to [38; 42] Hz using a hamming-windowed zero-phase forward and reverse fourth-order Butterworth infinite impulse-response (IIR) filter.

#### 3.5.4 Statistical design and analysis

To answer the research question of how cognitive tasks modulate the power and propagation of 40 Hz signals, we tested for systematic differences between the levels in the behavioural factors using non-parametric cluster-based statistics of the source reconstructed 40 Hz power [43]. The null-hypothesis for the visual attention experiment was that left and right attention trial conditions were exchangeable. In the arithmetic experiment, the null-hypothesis was that low and high difficulty trial conditions were exchangeable. For each of the three types of visual stimulation (0 Hz CON, 40 Hz ISF, 40 Hz LF), the exchangeability of the behavioural conditions was tested in a two-sided significance test using 1500 random permutations, exchanging paired condition averages within subject.

Contrasting experimental conditions within the same type of flicker stimulation will reveal the modulation effects of a given factor only if the assumption of system linearity is true, that is, we assume that the neural signal during the two experimental conditions are superimposed on a constant artifact at 40 Hz. In the case of non-linear coupling of the stimulators and the MEG system, the recorded neural signal may have an unpredictable interference with the artifact. In that case the artifacts cannot be guaranteed to cancel out, and the difference in neural signal between conditions may reflect a mixture of the modulation effect with the artifact.

#### 3.5.5 Exploratory control analysis

Given the results pertaining to the research questions (see sections 3.1 and 3.2), we decided to conduct exploratory analyses of the data to verify the methods described above (see exploratory results in section 3.3).

To verify that the third-order gradient rebalancing and PCA suppression of the 40 Hz EMI artifact worked as intended, we plotted the time series from one channel from 10 randomly selected trials in which 40 Hz stimulation took place. This was done for each participant before and after the artifact removal procedure. We plotted a time window from two seconds before stimulus onset and until the end of the trial to visually confirm that the EMI artifact arising from the stimulus onset was removed.

By plotting the PSDs for each stimulus condition and the contrast between 40 Hz and 0 Hz stimulus conditions, we verified that the 40 Hz signal had not been removed entirely in the process of de-noising the 40 Hz EMI artifact. With these, we confirmed that the spectrum was entirely aperiodic during stimulation with 0 Hz CON (except 50 Hz line noise), that the spectrum had a clear peak at 40 Hz during 40 Hz ISF/LF stimulation, and that the contrast between 40 Hz ISF/LF and 0 Hz CON revealed a spectrum with power of difference 0 dB for all frequencies except the 40 Hz bin.

As the contrasts between 40 Hz ISF/LF and 0 Hz CON do not eliminate the effects of the artifact, we also plotted the contrast spectra within the 40 Hz ISF and LF conditions. Between conditions with the same stimulus, the 40 Hz artifact is expected to be constant, and thus the difference should entirely remove the artifact, revealing only the modulation (interaction) effect of the behavioural task. Here, the spectrum is expected to be distributed around a power difference of 0 dB, with identical variance across frequencies, except for the 40 Hz bin which can be either offset from or distributed around 0 dB.

## 4 Discussion

Positive indications for 40 Hz treatment of AD continuously emerge [4–11, 16] along with plausible hypotheses for the mechanism of action involving glymphatic clearance [18, 21]. Traditionally used 40 Hz luminance flicker is associated with significant discomfort, compared to luminance-matched non-flickering lights [23]. To accommodate the need for human-centred visual 40 Hz stimulation, invisible spectral flicker was proposed [31].

Forty hertz ISF was found to be significantly less flickering and more comfortable than 40 Hz LF, though at the cost of a reduction in the evoked 40 Hz power.

As the promotion of glymphatic clearance from 40 Hz sensory stimulation [21] appears to be a local phenomenon controlled by the envelope of gamma oscillations [19], it is desirable to track the power and propagation of the 40 Hz signal.

Magnetoencephalography provides a good trade-off between temporal and spatial resolution for tracking the propagation of 40 Hz signal. However, 40 Hz ISF can not be produced with a standard RGB projector but requires specialised LED hardware that is not directly compatible with MEG.

The propagation of various types of invisible flicker has been studied using MEG by using a high-speed 1440 Hz projector, thus coining it *rapid invisible frequency tagging* (RIFT) [44–47]. These studies use RIFT for tagging and tracking certain cognitive mechanisms by embedding high-frequency luminance modulated targets in behavioural tasks, However, RIFT is considered invisible only for frequencies above 60 Hz [47], and as the projector is limited to spectra comprised only of red, green, and blue, it remains impossible for it to produce ISF.

Here, we took the first steps to use MEG to track the propagation of 40 Hz ISF. Our results as outlined below show that for now, this remains an unsolved challenge. We provide several suggestions resolving these seemingly incompatible modalities.

### 4.1 Behavioural effects

#### 4.1.1 Visual attention

In the visual attention experiment, only minor variations where observed in the behaviour. No significance tests were a-priori planned or performed, as the behavioural experiment that was part of the MEG recordings was not designed and powered for that purpose.

Despite this, we had expected the three types of visual stimuli to potentially influence behavioural responses. In the event of non-zero difference between the 40 Hz LF positive control and the 0 Hz CON negative stimuli on behaviour, we expected both negative and positive differences to be plausible. Either 40 Hz LF could reduce the response accuracy and increase the response time due to flicker as a distractor [48, 49], or inversely improve both due to increased focus of visual attention as it has been shown for 40 Hz binaural beats [50]. In extension thereof, we hypothesised that 40 Hz ISF could be closer to one or the other and in either direction. However, there was no indication for any differences in response time or accuracy between stimulation conditions. We did not expect differing behavioural responses between tasks with congruent and incongruent discrimination gratings, which was also the case.

The high response accuracy suggests that it was, perhaps, too easy to discern the orientation of the discrimination grating even without attending to the cued side, and as such there may not have been a benefit from attending to the cued side. Assuming this to be the case, the modulation effect of the lateralised attention on the 40 Hz evoked response from the brain would be smaller or might even be absent. To improve this for future experimental designs, the difficulty of the task would need to be better titrated, and/or the task would have to be modified to be able to assess a behavioural benefit.

#### 4.1.2 Arithmetic difficulty

As with the visual attention experiments, we hypothesised that the 40 Hz ISF and LF in the arithmetic experiment could have either a negative distractor-like influence on behavioural response time or accuracy, or a positive influence as it has been shown previously with 40 Hz binaural beats on working memory [51]. However, neither turned out to be the case as there were no notable differences in response time or accuracy between the three stimuli.

We observed shorter response times for trials where the presented answer was correct - though there was only a slightly higher response accuracy - compared to incorrect. This suggests that participants where ready to respond when they saw the correct answer, but had to re-evaluate when seeing a wrong answer.

Finally, we observed the expected difference in the response accuracy between low and high difficulties. The difficulty also appears to have affected the response time which was slightly shorter for the low difficulty. The observation that response accuracy for the low difficulty condition almost saturates toward the upper limit of 100% indicates that it required very little cognitive load, suggesting that the desired contrast in cognitive load was achieved.

### 4.2 MEG data

Unfortunately, the channel- and source-level MEG results did not offer neurophysiological insights as to whether cognitive tasks modulated the power or propagation of the 40 Hz stimulation input signal. Across spectral analysis, sensor-level topography, and source reconstruction, no meaningful inference could be made due to the uncontrolled mixture of 40 Hz neural signals and 40 Hz EMI artifacts.

The lack of an observable peak 40 Hz power in the contrasted PSDs in figs. 3 and 6 panels B and C could be due to the absence of modulation effects of behaviour on 40 Hz power and propagation. However, heteroscedasticity across frequencies with a clear reduction in variance at the 40 Hz bin reveals that the superposition of 40 Hz EMI artifact and 40 Hz neural responses may have been imperfect. In a linear system, given the consistent spatial and temporal nature of the stimulator emissions across behavioural conditions, these should cancel entirely and reveal the modulated neural signal in a contrast of behavioural conditions. The suppression of 40 Hz power was likely the result of unintended effects of the EMI coupling, leading to non-linear behaviour in the MEG sensors. The suspected non-linearities subsequently affected the three safeguards put in place to disentangle the artifact from the neural signals.

We attempted to remove the 40 Hz EMI artifact using synthetic third-order gradients. However, these operate entirely by projection of and subtraction of a linear combination of reference and sensor outputs [37]. Thus, they assume the system to be linear to allow for cancellation of the superposed neural and EMI signals. The subsequent preprocessing step of projecting and removing residual 40 Hz artifacts using principal components estimated from the reference gradiometers also assumed system linearity. While a lot of the artifact appears from fig. 8 to have been suppressed, it is impossible to infer how much of the residual 40 Hz power is attributed to EMI and how much is from cortical activity. This also renders the spectral and topographic representations shown in figs. 3 and 6 non-interpretable, as the contrasts between conditions assumes linearity in the sense that effects are additive across factors.

Adaptive beamforming was employed for the simultaneous purposes of investigating the propagation of the 40 Hz brain activity and further eliminating environmental noise. The LCMV beamformer aims to construct a series of spatial filters with a passband pointed at each (discretised) location in the brain and stopband everywhere else [38]. However, this requires orthogonal sources, and while LCMV is robust to moderate correlation between spatially distinct sources, it is intended to suppresses correlated sources [38]. In anticipation of symmetric lateral time-locked evoked responses, we applied a symmetry constraint for the LCMV beamformer to fit double-dipoles symmetric across the midline [42].

Despite these attempts, we found no systematically different sources between lateralisation task levels in the visual attention experiment. In the arithmetic experiment, between the three types of visual stimuli, we found three near-identical sources with wide support in the arithmetic difficulty contrast. We did not have any neurophysiological interpretation of these results, in which both the 0 Hz CON negative control and 40 Hz LF positive control appeared to result in the same degree of modulated 40 Hz source power from the task.

The double-dipole is spatially specific, and thus its usefulness is limited to correlated sources defined by discrete pairs when defining the grid. Thus, it does not account for correlation between sources in the brain and unmodelled sources outside the brain.

Since the two stimulators were operating independently of each other and thus had an uncontrolled and unknown phase difference between them, the chance of neural 40 Hz sources being at least partially correlated with either of them is fairly high. The EMI power was likely orders of magnitudes higher than any neural signals, which were most likely cancelled by the beamformer.

### 4.3 Future modifications to experimental design

The suspected non-linear behaviour of the MEG sensors as a consequence of 40 Hz EMI introduced uncontrollable confounds and potential suppression of the outcome measure, and thus the experimental setup needs to be reconsidered. Here, we provide several avenues of improvement that might allow for the use of MEG to track propagation of 40 Hz ISF.

#### 4.3.1 Hardware and setup

We suggest removing the 40 Hz EMI artifact entirely, rather than attempting to control it by means of signal processing and statistical designs that require system linearity.

Although challenging, this could be achieved by moving the light sources from inside (see fig. 11) to outside the MSR. The delivery of spectrally specific light stimulation to the MSR is limited to the use of the specific LED chips in the stimulators. Conceivably this could be be achieved by two different approaches: 1) using lenses and mirrors to focus and direct the light from the LED along the projector projection path (fig. 15 panel A), or 2) by employing fibre-optic coupling of the light directly from the LED chip using fibres fed through the MSR wall (fig. 15 panel B). Either would allow both the LEDs and their current-supplying drivers to remain fully outside the MSR.

**Fig 15.**
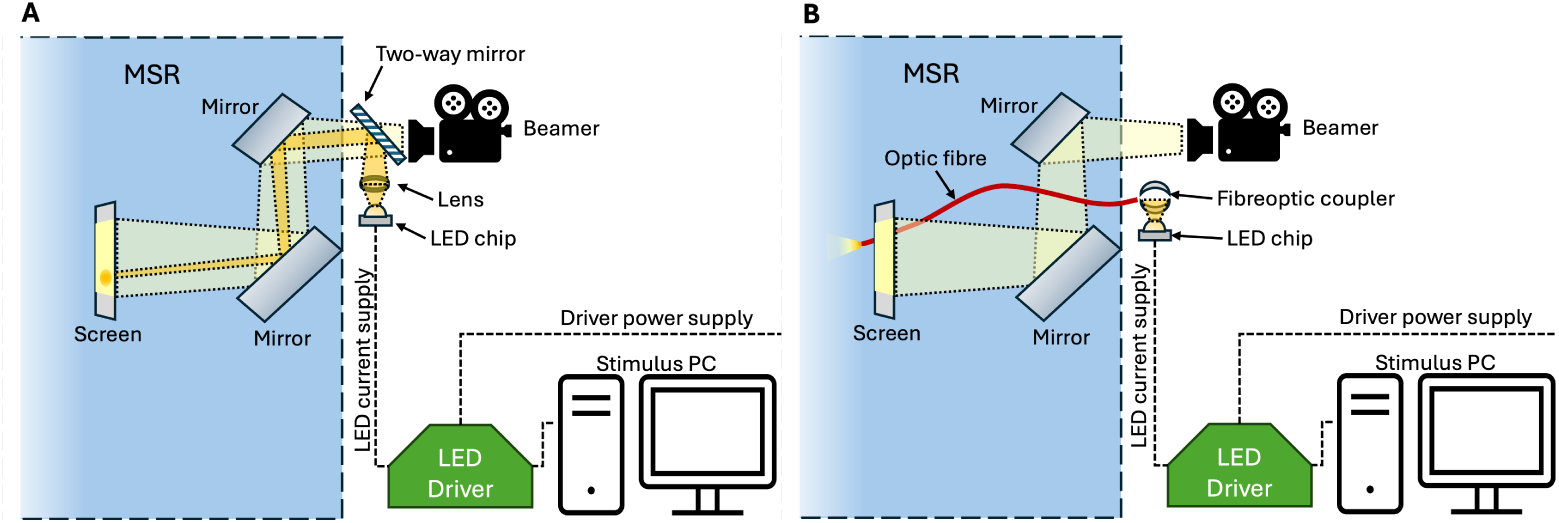
Alternative light-delivery setup: The 40 Hz LED light sources can feasibly deliver stimulation by two proposed setups. **A)** A geometrically delicate optic pathway, focusing and reflecting the LEDs outside to the screen the MSR could be a fibre-free solution. **B)** Though optic fibres, coupling the LED light output outside the MSR, the light can be pointed at the screen or directly towards the participant. Both implementations would suffer from reduced illumination, and thus their ability to evoke a reliable 40 Hz response must be confirmed.

The lens-based approach is geometrically challenging in terms of consistently focusing the light output to the correct spot on the presentation screen. The fibre-optic approach might practically be easier and has been implemented previously with 40 Hz ISF in an fMRI study [32]. The light-emitting terminals of the fibre-optic cables could be positioned closer to the participant than the projection screen with the task instructions from the regular projector, and thus it may require a redesign of the participant’s interaction with the two in a visual attention experiment.

However, both proposed setups will suffer from reduced illumination and may distort the matching of the luminance of the two alternating LED sets in 40 Hz ISF, which should therefore be re-calibrated according to the procedure specified in Section 2.2 of [31].

#### 4.3.2 Design of experiments

The statistical power of the studied effects is dependent on the number of repetitions and participants, effect magnitudes, and the within- and between subjects variance [52]. Thus, there is motivation to improve all of these.

The number of participants was primarily limited by time and budget as practical constraints, but could be increased. The number of repetitions is not easily increased under the current paradigm, as multiple participants reported getting tired towards the end of data collection. Instead, the number of 40 Hz ISF and LF repetitions can be increased by 50% by eliminating the 0 Hz CON negative control, testing that condition only in the verification procedure (see sections 5.4.1 and 5.4.3). The returns are, however, expected to be diminishing from increasing repetitions and participants [52].

The observation that one subject answered randomly in all conditions of the visual attention task, and another answered randomly in two specific conditions, indicates that the experiment itself can be improved, and the purpose of the tasks is to be explained better and trained more. This might improve both the effect magnitudes and reduce the within- and between subject variance. In a future design, there should be a benefit associated with correctly maintaining the gaze fixed on the centre of the presentation screen, but attending to the cued light stimulus. This could be achieved by making the task more difficult, and thus requiring more engagement with both.

Finally, given that the 40 Hz EMI artifact can be pre-emtively eliminated by placing the LEDs outside the MSR, tracking of the 40 Hz signal does not have to rely entirely on behavioural modulation of the power and propagation. Instead, it can be investigated with stronger conditional contrasts such as the comparison of 40 Hz ISF and 0 Hz CON directly during a task-based or task-free experiment.

### 4.4 Verification procedure

Here, we propose a set of experimental checks to verify both the hardware setup and the experimental design. All of the following should be performed in the MSR during MEG recording prior to data collection.

#### 4.4.1 Absence of 40 Hz EMI

With the goal of eliminating the 40 Hz EMI artifact by removing the light sources from the MSR, it will be necessary to verify that no 40 Hz coupling takes place in an empty room with the light sources on. This can be achieved by conducting an empty room recording with the light sources (however delivered) first off, and then on each of the 0 Hz Con, 40 Hz ISF, and 40 Hz LF settings. A spectral analysis during each of these three recordings should confirm the absence of a peak in power at 40 Hz.

#### 4.4.2 Presence of 40 Hz evoked response to light stimulus

It is necessary to verify that the delivered 40 Hz modulated light (however realised) can evoke a 40 Hz response. If the absence of 40 Hz EMI is confirmed, this can be achieved by stimulating a pilot participant with 40 Hz ISF and 40 Hz LF in a task-free setting. Time-frequency analysis should reveal a steady state visual evoked response at 40 Hz, starting from stimulus onset.

#### 4.4.3 Modulation of 40 Hz by cognitive load

Our behavioural paradigms were inspired by those used in [33]. They were modified to allow us to estimate the modulation of cognitive load on 40 Hz power and propagation between task levels instead of comparing the task to task-free setting. Although this is a truer representation of the modulation effect, it requires a formal verification of similar modulation effects. This can be achieved by conducting the the visual attention and arithmetic experiments exclusively with the 0 Hz CON negative control and 40 Hz LF positive control light stimulation. Without 40 Hz ISF, the need for specialised hardware is eliminated and the light stimulation can be delivered with the high-frequency RGB projector as it has been done in other high-frequency flicker experiments [45–47]. Ideally, the luminance of the light stimuli delivered by the projector should be matched to those of the specialised hardware.

## 5 Conclusion

The necessity for specialised electronics hardware to stimulate using 40 Hz invisible spectral flicker makes tracking of the cortical propagation using magnetoencephalography (MEG) currently still an unsolved challenge.

Under the assumption of constant noise in a linear system, we expected that the confounding 40 Hz electromagnetic interference from the visual stimulator could be disentangled from the cortical 40 Hz response. We attempted this by four pre-planned safeguards: 1) customisation of visual stimulator for improved electromagnetic compatibility, 2) linear transformations for active noise cancelling in MEG preprocessing and de-noising, 3) source reconstruction of the neural source using adaptive beamformer filtering for suppression of external sources, and 4) contrasted behavioural conditions for modulating the 40 Hz neural power and propagation.

The planned efforts, however, were impeded by what we suspect to be non-linear behaviour of the MEG sensors during 40 Hz stimulation. This rendered the mixture of artifact and neural signals uncontrolled and confounded, thus making inference about the behavioural modulation of 40 Hz power and propagation impossible.

We present a set of suggestions for improvement through further hardware modifications and experimental design for behavioural tasks optimisation, as well as a verification procedure for its feasibility. These may render the future use of MEG for tracking 40 Hz invisible spectral flicker possible.

## Acknowledgments

The authors would like to thank Donders Centre for Cognitive Neuroimaging for providing the MEG and MRI imaging facilities, OptoCeutics for providing the brain stimulation devices, Gustavo Feijóo Carrillo and Uriel Plönes for technical assistance in customising the brain stimulation devices, Miranda Naaktgeboren for MEG training, and Paul Gaalman for MRI assistance.

## Notes

### Competing Interest Statement

M.H. is employed as an industrial Ph.D. student by OptoCeutics ApS, funded partially by the Innovation Fund Denmark (case no.: 1044-00177B), but enrolled as a Ph.D. student under the supervision of K.M. at the Technical University of Denmark (DTU), and has stock options in OptoCeutics ApS.
E.S. declares no conflict of interest.
H.H. is a former employee of and owns shares in the company OptoCeutics ApS.
M.C. is chief scientific officer of and owns shares in the company OptoCeutics ApS.
K.M. declares no conflict of interest.
R.O. declares no conflict of interest.
The costs of visual stimulation equipment and modifications thereof are covered by OptoCeutics ApS.

